# CFL1-dependency of discrete cell behaviours driving mouse spinal zippering

**DOI:** 10.1101/2023.11.07.565942

**Authors:** Abigail R Marshall, Andrea Krstevski, Henry Crosswell, Rahul Shah, Eirini Maniou, Nicholas DE Greene, Andrew J Copp, Gabriel L Galea

## Abstract

Epithelial fusion is critical for formation of many embryonic tissues. The developing mammalian spinal cord fuses by zippering, through cell behaviours dependent on F-actin turnover. We propose a caudal-to-rostral sequence of surface ectoderm cell behaviours which drive zippering progression in mice, and test the requirement for the F-actin severing protein CFL1 in this process. Key zipper-advancing behaviours are *i.* constriction of supracellular actomyosin cables; *ii.* extension of short-lived (<2 min) filopodial or more persistent lamellipodial protrusions, which we live-image establishing ‘pioneering’ contacts across the midline; remodelling of *iii.* cell-cell and *iv.* cell-ECM adhesions; *v.* zipper advancement through constriction of leading-edge cell borders. In wildtype embryos CFL1 is enriched at the leading edge of surface ectoderm cells adjacent to the zippering point and localised within some protrusions. Conditional *Cfl1* deletion in the surface ectoderm produces more prominent F-actin stress fibres, makes filopodial protrusions excessively stable, diminishes their exploratory movements which may establish nascent contacts, and impairs junctional constriction of cells at the point of fusion. Zippering speed is reduced ∼30% in conditional *Cfl1*-deleted embryos and ∼30% develop spina bifida. We therefore propose that impaired filopodial dynamicity limits the sequence of cell behaviours driving spinal zippering in mice, predisposing to spina bifida.

## Introduction

Epithelial fusion is crucial for morphogenesis of many tissues including the palate, urethra, optic fissure, body wall and central nervous system^1^. Some fusion events are relatively well described as a sequence of discrete cell behaviours. For example, *Drosophila* embryos undergo dorsal closure through a ‘zippering’ mechanism coordinated by actomyosin purse-strings overlying contraction of the amnioserosa^2,3^. In contrast, they close their cardiac vessel through a ‘buttoning’ process requiring stabilisation of homophilic cell-cell adhesions on filopodial protrusions^4–6^. These two modes of epithelial fusion by zippering versus buttoning can be employed by different species to achieve equivalent morphogenetic events: as exemplified by closure of the spinal neural tube in mouse versus chicken embryos^7–9^. The neural tube is the embryonic precursor of the brain and spinal cord which closes from a flat neural plate into a quasi-cylindrical tube. Bending of the neuroepithelium brings the left and right halves (neural folds) into apposition at the dorsal midline, where the surface ectoderm (SE, future epidermis) fuses to advance closure^7,10,11^. Whereas in chickens the spinal SE meets at interspersed buttoning points^8,12^, mouse and human^13,14^ embryos zipper unidirectionally from the hindbrain-cervical boundary to the low-spine. Interruption of zippering progression along the ∼3.5 mm length of the mouse embryonic axis – due to SE-intrinsic defects or secondary impediments such as unfavourable tissue biomechanics – causes spina bifida^9–11,15,16^.

The cellular and molecular bases of neural tube zippering have been elegantly delineated in the hemichordate *Ciona*^17,18^. In these embryos, epidermal cells which meet at the point of epithelial fusion (‘zippering point’) establish new cadherin-dependant contacts and enrich myosin at their leading edge, causing sequential cell border constriction as they leave the neural fold rim^17,18^. We and others have identified conserved elements of this sequence during zippering of the mouse spinal neural tube – the open region of which is called the ‘posterior neuropore’ (PNP). For example, mouse zippering point SE cells have similarly constricted leading-edge borders, forming a semi-rosette-like organisation^19,20^. The pair of SE cells directly at the embryonic midline have the most constricted cell borders, whereas those further away along the neuropore rim are less constricted^19^. Semi-rosette border constriction requires cell-extracellular matrix (ECM) adhesion: we previously reported that conditional deletion of integrin β1 (*Itgb1*) in the SE diminishes semi-rosette border constriction and stalls zippering in the low-spine, causing spina bifida in 56% of embryos^19^. Whereas *Ciona* embryos only enrich myosin in the cells directly at the zippering point^17^, mouse SE cells assemble supracellular actomyosin cables which link cells >200 µm along the length of the neuropore^21^. These cables generate a tensile force which helps advance zippering^22,23^. Cable tension requires actomyosin constriction downstream of Rho/ROCK signalling^23^, and is transmitted between neighbouring cells through adherens junctions^21,24^.

SE cells at the neuropore rim also extend filopodial and lamellipodial protrusions reminiscent of those used by migrating mesenchymal cells^10,20,25^. Filopodial protrusions are formed under the control of the Rho-GTPase CDC42 at early zippering stages, approximately equating to the cervical and high thoracic spine^10^. These subsequently intermingle with lamellipodial-type protrusions formed under the control of an alternative Rho-GTPase, RAC1^10^. Lamellipodial protrusions replace filopodial ones by the level of the low-lumbar spine^10^. Conditional deletion of RAC1 eliminates lamellipodia, replacing them with filopodia in low-spinal zippering, but these are insufficient to sustain closure, producing spina bifida in ∼90% of embryos^10^. The necessity of filopodial protrusions for early spinal zippering is less clearly demonstrated because equivalent conditional deletion of CDC42 causes precocious extension of lamellipodia and severe embryo stunting^10^.

Despite protrusions and supracellular cables co-existing in the same SE leading edge, their regulators CDC42/RAC1 and Rho/ROCK act antagonistically to each other in other contexts^26–30^. CDC42, RAC1 and ROCK all inhibit cofilin (CFL)1, which binds and severs F-actin^31–34^. F-actin turnover is required for all cell behaviours involved in spinal zippering. It is polymerised through the action of formin and ARP2/3 complexes, stabilised by various binding proteins, and destabilised by severing proteins including CFL1 and destrin (also called actin depolymerising factor, ADF) ^35,36^. Mice globally lacking destrin are viable^37^, whereas global loss of *Cfl1* causes early embryo lethality with fully penetrant neural tube defects^38–40^, suggesting non-redundancy with destrin in this process. CFL1’s roles in the intricate interplay between F-actin polymerisation and severing have largely been studied using *in vitro* reactions or cultured cells^41–43^, particularly in the context of cellular extension of lamellipodia and filopodia during cell migration^44–47^. The relevance of these processes *in vivo* is evident in tractable model organisms and morphogenetic events, such as *Drosophila* dorsal closure^48^. Genetic variants of actomyosin regulators including CFL1^31^ have also been implicated in human neural tube defects. However, analysis of their functions during mammalian morphogenesis has been hindered by redundancy^49^, and early embryonic lethality when non-redundant genes are disrupted^10^.

CFL1 predominantly interacts with ADP-bound, older F-actin filaments, altering their configuration^50^ and recruiting actin interacting protein (AIP)1^51^, promoting severing of the filament. Actin binding is diminished by CFL1 phosphorylation, for example by LIM kinase (LIMK) activated downstream of Rho/ROCK signalling^31^. LIMK1 activity can also be increased downstream of CDC42 and RAC1, likely indirectly through inhibition of its phosphatase, slingshot^32–34^. CFL1 association with F-actin is also antagonised by binding to phosphatidylinositol 4,5-bisphosphate (PIP2), particularly in cell sub-domains close to the membrane such as in protrusions^31,52,53^. These multiple interactions with known regulators of mammalian spinal fusion make CFL1 a high priority candidate to regulate SE zippering behaviours, yet previous analyses of its functions have focused on the neuroepithelium^38,39^.

Here, we define a sequence of cellular behaviours hypothesised to drive spinal zippering in mice and use conditional gene deletion approaches to mechanistically dissect key cellular functions of CFL1 in this process. We demonstrate that CFL1 is enriched adjacent to the zippering point at early stages of spinal zippering, when filopodial protrusions predominate. Its regulation involves both differential sub-cellular localisation and tissue-level patterns of phosphorylation within the SE. Our systematic analysis of embryos conditionally lacking its expression in the SE demonstrates partially penetrant spina bifida and a ∼30% reduction in zippering speed associated with diminished dynamicity of filopodial zippering protrusions.

## Results

### Description of cellular behaviours underlying mouse spinal zippering

The mouse spinal zippering point is morphologically identifiable as a V-shaped meeting of the left and right sides of the PNP at its rostral extremity (Figure 1a). SE cells at the rim of the neural folds assemble supracellular cables and extend F-actin-rich zippering protrusions (Figure 1b). ECM is sparse around the neuropore rim^19^, but fibronectin fibrils are identifiable directly at the zippering point (Figure 1c). Here, cells are interlinked by tight junctions (Figure 1c) as well as by E-cadherin adherens junctions (Figure 1d). Adherens junctions punctuate the actomyosin cables at the neuropore rim, interlinking adjacent cells (Figure 1d). Cell borders engaged in the actomyosin cables lack leading edge E-cadherin, but progressively contract as they approach the zippering point (Figure 1d). They establish new junctions around their perimeter when they leave the zippering point (Figure 1d). These zipper-advancing behaviours can be described in a caudal-to-rostral spatial sequence: *i.* SE cable constriction, *ii.* extension of protrusions, *iii.* cell-cell and *iv.* cell-ECM adhesion, followed by *v.* leading edge border contraction advancing the zipper caudally. Each of these steps could be regulated by CFL1.

**Figure 1:**
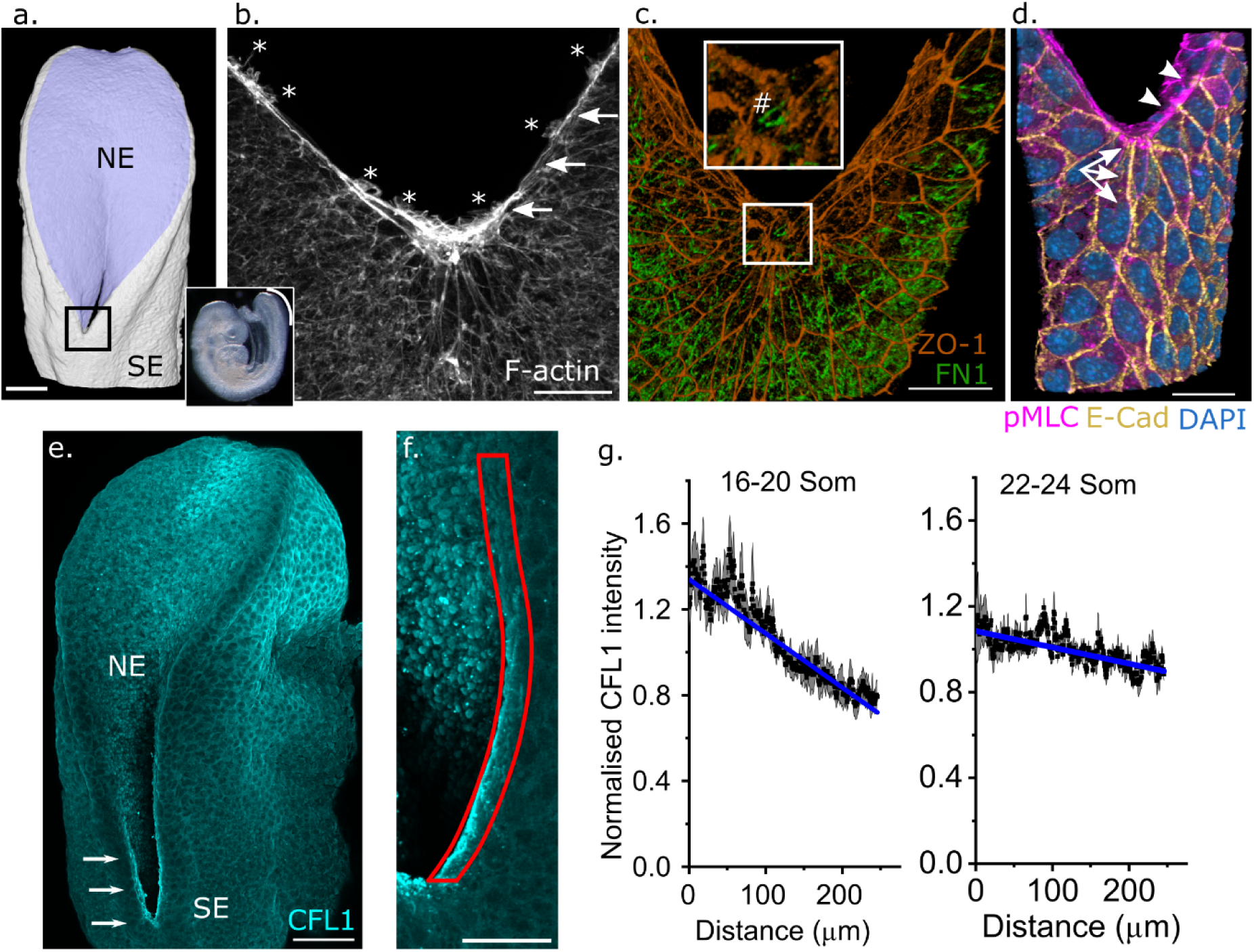
SE process and CFL1 localisation in spinal neural tube zippering. **a.** 3D reconstruction of an illustrative PNP indicating the neuroepithelium (NE) and SE (SE). The PNP is located under the white line shown in the brightfield image of an E9 mouse embryo. The black box indicates the zippering point. **b-d.** High-resolution confocal images of the zippering point. **b.** F-actin staining showing supracellular cables (arrows) and SE protrusions (*). **c.** ZO-1-labelled apical cell borders and fibronectin (FN1), illustrating its presence at the zippering point (#). **d.** Phosphorylated myosin light chain 2 (pMLC), E-cadherin and nuclear labelling showing SE midline adherens junctions (arrows) and junctions linking cells forming the supracellular actomyosin cables (arrowheads). Scale bars = 25 µm. **e-f.** Wholemount confocal images showing CFL1 expression. **e.** 11 somite embryo with CFL1 enriched at the zippering point and along the neuropore margin (arrows). Scale bar = 100 µm. **f.** 18 somite embryo with the red boundary indicating the region in which CFL1 intensity was quantified in **g**. Scale bars = 50 µm. **g.** Quantification of CFL1 intensity along the neuropore rim in embryos with the indicated somite stage groupings. 0 µm is the zippering point. Points represent the average ± SEM CFL1 intensity from six embryos per somite grouping, with the average intensity of each embryo normalised to 1. The slopes of the two regressions are significantly different from each other: F-test p < 0.001.

### CFL1 regulation by phosphorylation and sub-cellular localisation during spinal zippering

CFL1 is expressed throughout the SE and neuroepithelium of the PNP (Figure 1e) but is enriched at the leading edge of SE cells adjacent to the zippering point (Figure 1e-g). The enrichment is developmental stage specific – becoming less pronounced in embryos at more advanced somite stages (Figure 1g). This suggests CFL1 is targeted to the leading edge of SE cells bordering the neural folds, where they form F-actin cables and extend protrusions (Figure 1b), at early stages of PNP closure.

High-resolution imaging immunolocalises CFL1 within some of these protrusions, but less so in others (Figure 2a). Similarly phosphorylated CFL1 is also present in some protrusions but not others (Figure 2b). Beyond sub-cellular localisation and phosphorylation, we also observe whole-cell and tissue-level patterns of CFL1 regulation. Our adaptive ‘surface subtraction’ image processing method^16^ allows selective visualisation of the SE. This reveals heterogenous pCFL1 abundance throughout the cytoplasm of adjacent SE cells and marked reduction of pCFL1 immunolocalization in mitotic cells of both the SE and neuroepithelium (Figure S1a). pCFL1 is particularly enriched in midline SE cells which have completed zippering (Figure 2c,e, S1b). Pharmacological inhibition of ROCK globally diminishes pCFL1 intensity (Figure 1c-d), suggesting regional or cell-level activation of this pathway contributes to the observed pattern of CFL1 regulation.

**Figure 2:**
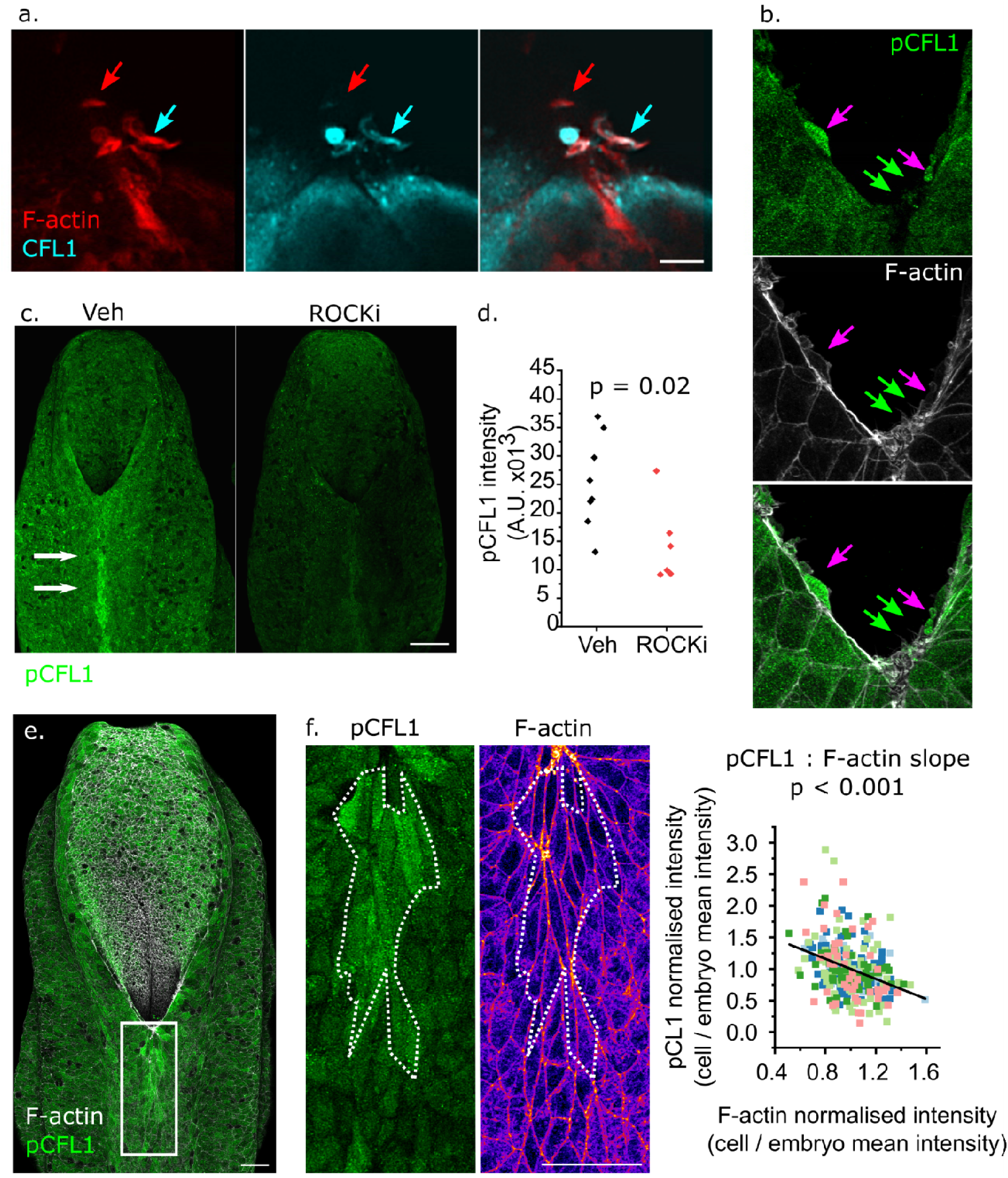
SE CFL1 regulation by subcellular localisation and cell-level phosphorylation. **a.** AiryScan super-resolution confocal image showing SE F-actin-rich protrusions with high (cyan arrow) or low (red arrow) level of CFL1 localisation. Scale bar = 5 µm. **b.** Surface-subtracted AiryScan image showing F-actin rich protrusions with (magenta arrows) and without (green arrows) phosphorylated CFL1. Scale bar = 5 µm. **c.** Surface-subtracted wholemount images of pCFL1 in a vehicle (Veh) control and ROCK inhibitor (ROCKi, 10 µM for 4 hours) treated embryo. Arrows indicate a stripe of midline SE cells with highly phosphorylated CFL1. Scale bar = 100 µm. **d.** Quantification of pCFL1 staining intensity in the dorsal SE of vehicle and ROCK-inhibited embryos. Points represent individual embryos. **e.** Wholemount image showing colocalization of pCFL1 and F-actin. The white box indicates the region shown in ‘**f**’. Scale bar = 50 µm. **f.** High resolution images of the dorsal SE pCFL1 and F-actin (shown in Fire LUT to help visualise differences in intensity). Scale bar = 50 µm. The dashed white border indicates a region of several cells with high pCFL and low overall F-actin staining. The graph quantifies pCFL1 and F-actin intensity. Both pCFL1 and F-actin were independently normalised to 1 in each embryo (cell intensity / embryo average intensity) so that cells from different embryos can be compared. Points represent individual cells, colours indicate different embryos (293 cells from 5 embryos). P value for the slope based on Pearson’s regression.

We hypothesised that cell-to-cell heterogeneity of pCFL1 levels may reflect differential requirement for F-actin turnover. F-actin is brightly localised at cell-cell junctions, and forms stress fibre-like formations of variable intensity between adjacent cells (Figure 2f). Quantification of cell-level pCFL1 and F-actin mean intensity in SE cells along the embryonic midline shows a highly significant negative correlation between them (Figure 2f), which was unexpected given that knockout of CFL1 causes F-actin accumulation^38^. This nonetheless provides evidence of cell-level association between CFL1 inhibition and F-actin in the SE.

### CFL1 in the early SE enhances zippering speed

To test the functional consequences of loss of *Cfl1*, we initially deleted it using *Cdx2^Cre^* (henceforth called cCre). This Cre-driver recombines in the spinal neuroepithelium from early stages of neural tube closure, with progressive expansion of its recombination domain to the SE and mesoderm around the PNP at later stages (Figure 3a). Consequently, cCre initially deletes CFL1 in the neural tube (Figure 3b), before globally supressing its expression at later stages (Figure 3c). Surprisingly, this multi-tissue deletion of *Cfl1* does not produce marked neural tube phenotypes, with only a minority of conditional knockout embryos developing a kinked tail abnormality (Figure 3d).

**Figure 3:**
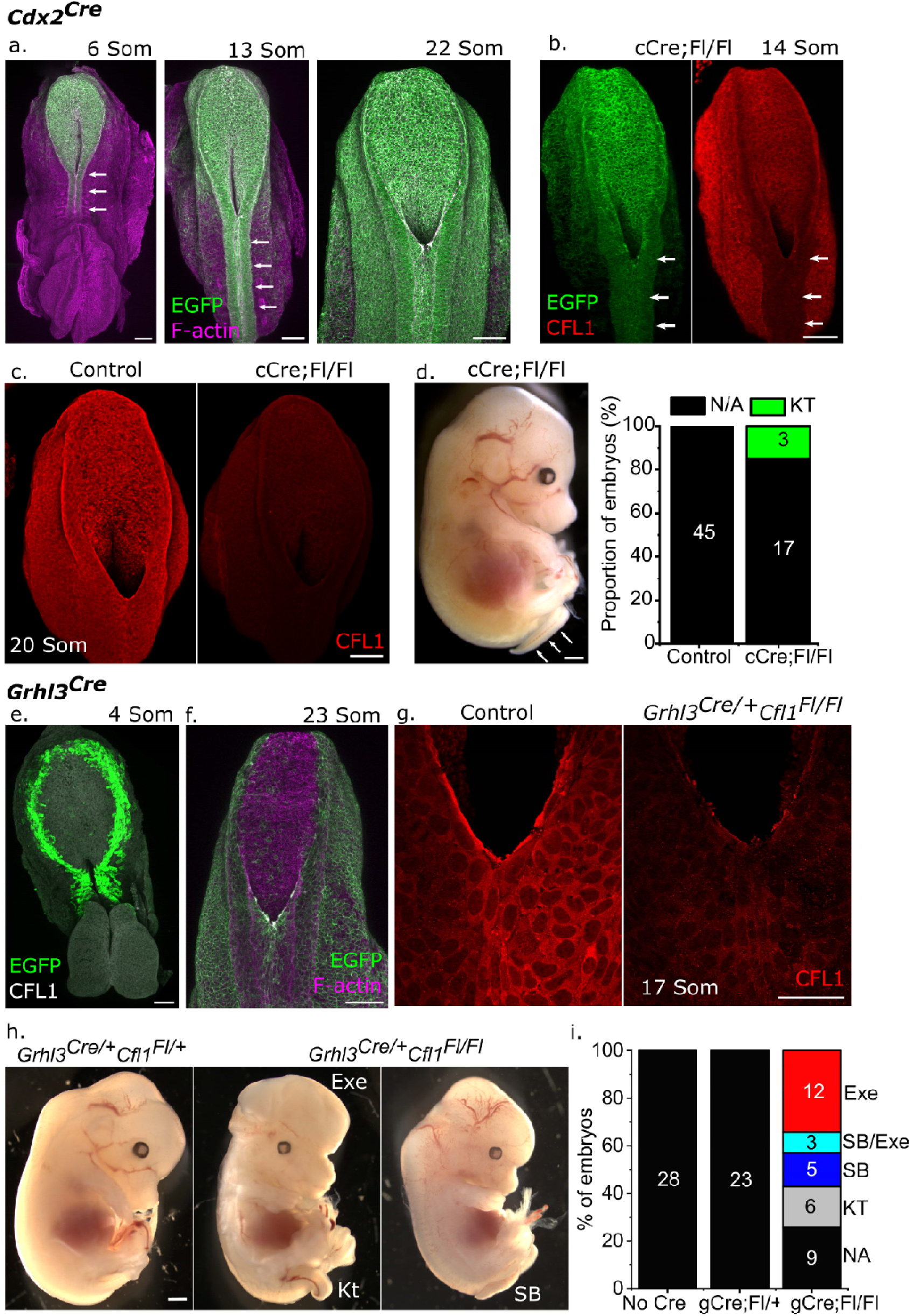
Cfl1 in the early SE contributes to neural tube closure. **a.** Pattern of Cdx2^Cre^ recombination (EGFP) visualised using the mTmG reporter in embryos of the indicated somite stages. Note predominant recombination in the neural tube (arrows) in the earlier embryos, extending to the SE and mesoderm at later stages. **b.** Wholemount image showing endogenous EGFP (mTmG reporter) and CFL1 immunolocalisation in a Cdx2^Cre/+^Cfl1^Fl/Fl^ (cCre;Fl/Fl) embryo with 14 somites. Note CFL1 loss in the cCre recombination domain (e.g. neural tube, arrows). **c.** Immunofluorescence comparison of CFL1 immunolocalisation in a control and littermate cgCre;Fl/Fl embryo with 20 somites. **d.** E14.5 cCre;Fl/Fl with kinked tail and quantification of the proportion of phenotypes observed. Numbers indicate the number of fetuses with each phenotype. **e-f.** Wholemount image showing endogenous EGFP (mTmG reporter) in Grhl3^Cre/+^ embryos with (**e**) 4 somites and (**f**) 23 somites. **g.** Immunofluorescence comparison of CFL1 immunolocalisation at the zippering point in a control and littermate Grhl3^Cre/+^Cfl1^Fl/Fl^ (gCre;Fl/Fl) embryo with 17 somites. **h.** Brightfield images showing kinked tail (KT), exencephaly (Exe) and spina bifida (SB) in gCre;Fl/Fl fetuses. **i.** Quantification of the proportion of phenotypes observed in gCre;Fl/Fl and littermate control fetuses. Numbers indicate the number of fetuses with each phenotype. NA = no abnormality, SB/Exe = combined SB and Exe in the same fetus. All SB fetuses also had Kt. Scale bars = 100 µm except d,h = 500 µm.

Use of cCre largely fails to inactivate CFL1 in the early SE, where we observed preferential CFL1 localisation adjacent to the zippering point (Figure 1e). An alternative Cre drive, *Grhl3^Cre^*, recombines in the early SE even before the initiation of neural tube closure (Figure 3e), eventually recombining the entire SE around the PNP (Figure 3f) and diminishing CFL1 at the zippering point in conditional knockout embryos (Figure 3g). This earlier deletion of *Cfl1* from the SE produces severe neural tube defects including partially-penetrant failure of cranial closure (exencephaly, not investigated further here) and spina bifida (Figure 3h-i). Thus, CFL1 in the early SE promotes completion of spinal neural tube closure.

Consistent with a SE-specific early role of CFL1, PNP length is significantly longer in *Grhl3^Cre/+^Cfl1^Fl/Fl^* (henceforth gCre;Fl/Fl) embryos than littermate controls by the 15-18 somite stage range (Figure 4a). To directly compare zippering speed, we injected DiI landmarks adjacent to the zippering point of cultured embryos and quantified their progression of zippering after culture (Figure 4b). gCre;Fl/Fl embryos zippered ∼30% slower than controls (Figure 4c). To begin exploring the basis for this zippering deficit, we first tested general cellular consequences of CFL1 loss in the SE. These cells normally have short stress fibre-like F-actin filaments below their apical cell-cell junctions (Figure 4e). gCre;Fl/Fl embryos still assemble equivalent junctional F-actin but have visibly longer and more prominent stress fibres (Figure 4e), consistent with the known roles of CFL1 in severing F-actin filaments *in vitro*^54^. Loss of CFL1 does not alter SE mitotic index (Figure S2a-b), but results in larger aggregates of apoptotic bodies (labelled with cleaved caspase 3) along the embryonic midline, where smaller apoptotic aggregates can also be observed in control embryos (Figure S2a). Apoptotic aggregates were not observed around the zippering point in either genotype.

**Figure 4:**
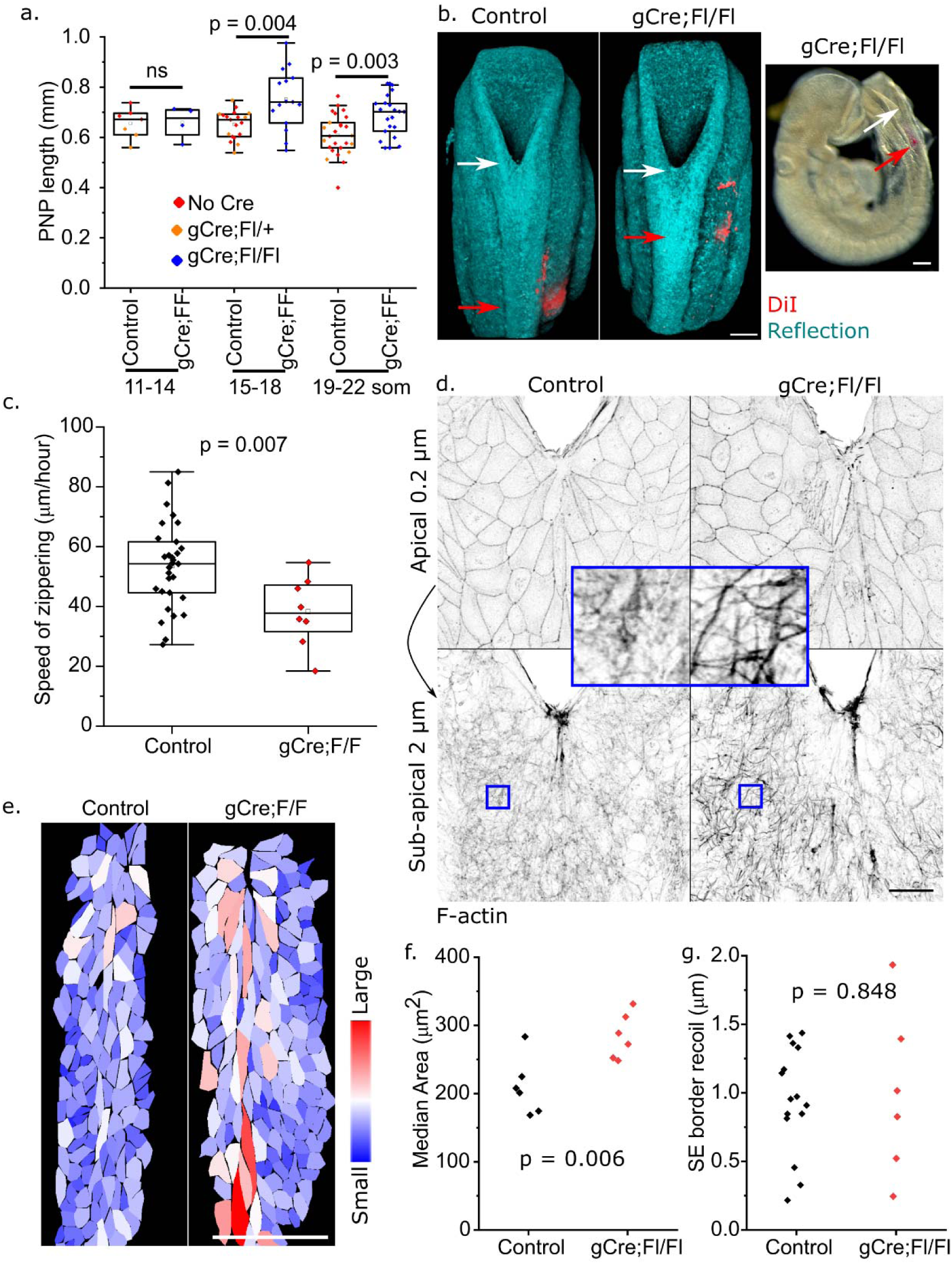
Cfl1 deletion in the SE impairs zippering, F-actin and cell size regulation, but not tension. **a.** Quantification of PNP lengths in embryos within the indicated genotypes and somite ranges. gCre;Fl/+ embryos are indistinguishable from Cre-negative littermates and are considered controls henceforth. **b.** Visualisation of zippering speed quantification using DiI landmark injection (red arrow) and confocal reflection imaging to quantify distance to the zippering point (white arrow) after 6 hours in culture. **c.** Comparison of zippering speed between E9 littermate control and gCre;Fl/Fl embryos (approximately 14-18 som at the start of culture). **d.** AiryScan visualisation of F-actin in the SE of a control and gCre;Fl/Fl embryo with 18 somites. The Z-stacks were sub-sampled using our adaptive surface subtraction method to only visualise the apical 0.2 µm in the top panel (showing F-actin at cellular junctions in both genotypes) and sub-apical 2 µm below it, visualising F-actin stress fibres within the cell body. Blue boxes indicate the positions magnified in the insert. **e-f.** Heatmap representation (**e**) and quantification (**f**) of projected SE cell areas along the dorsal midline immediately rostral to the zippering point in control and gCre;Fl/Fl embryos. Points represent the median cell area for individual embryos. **g.** Quantification of recoil following laser ablation in midline SE cell borders of control and gCre;Fl/Fl embryos. Points represent individual embryos.

Dorsal midline SE cells from gCre;Fl/Fl embryos are significantly larger than in controls (Figure 4e-f). We tested whether this increase in cell area is due to mechanical stretching, by using laser ablation to infer tension at cell-cell junctions rostral to the zippering point. Junctional tension was equivalent between control and gCre;Fl/Fl embryos (Figure 4g). Consistent with this, nuclear localisation of the mechanoresponsive transcription factor YAP^55^ was also equivalent between groups (Figure S2c-d). Thus, loss of SE CFL1 causes anticipated changes in F-actin organisation and quantifiable changes in cell shape, without directly impacting readouts of epithelial biomechanics.

### Analysis of zippering steps identifies CFL1-facilitated processes

In order to pinpoint deficiencies which may account for the reduction in zippering speed of gCre;Fl/Fl embryos, we analysed the spatial sequence of cell behaviours proposed to drive spinal zippering (Figure 5a), starting with the SE actomyosin cables. These are equivalently enriched for phosphorylated myosin in control and gCre;Fl/Fl embryos (Figure 5b). We used laser ablation to test the tension generated by these cables (Figure 5c), confirming that cable tension is not diminished in gCre;Fl/Fl embryos (Figure 5d). Cable-producing cells also extend zippering protrusions (Figure 5a’). While the density of lamellipodial protrusions around the zippering point was not significantly different between genotypes (Figure S3a), filopodial density is significantly greater in gCre;Fl/Fl than control embryos (Figure 5e-f). Exuberant filopodial protrusions persist even over the fused portion of neural tube in at least a subset of gCre;Fl/Fl embryos (or dynamically, so as to only be observed in some fixed embryos), producing an expanded area of protrusive activity rostral to the zippering point (Figure S3b-c).

**Figure 5:**
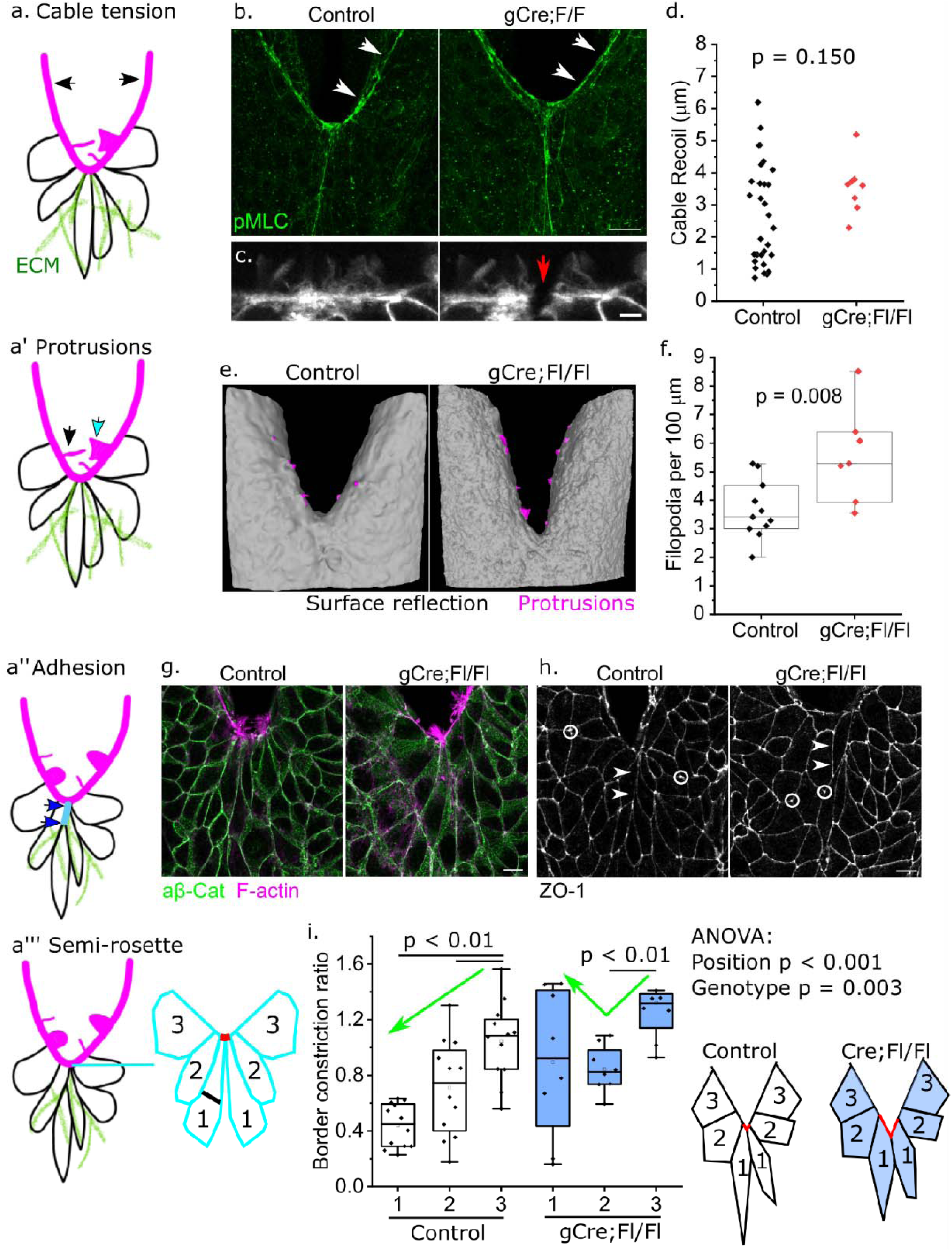
Systematic analysis of the zippering sequence in Grhl3^Cre/+^Cfl1^Fl/Fl^ embryos. **a-a’’’.** Schematic illustration of proposed sequence of zippering cell behaviours. a. Arrows indicating the SE supracellular cables (functionally analysed in b-d). **a’** Cyan arrow indicates lamellipodial (see Figure S3a) and black arrow indicates filopodial protrusions (analysed in e-f). **a’’** Blue arrows indicate new cell-cell junctions (visualised in g-h) with underlying cell-ECM adhesions (see Figure S3c). **a’’’** Schematisation of zippering point cell organisation in a semi-rosette (analysed in i). Cells are numbered in left/right pairs with 1 at the mid-line in contact with the most rostral point of the zippering point. Constriction of the cable-edge cell border is calculated as the ratio of the free border length (red lines) divided by the width of the cell body at half its length (black line). **b.** Immunolocalisation of pMLC in control and gCre;Fl/Fl littermate showing equivalent localisation in both. Scale bar = 25 µm. **c-d.** Visualisation (c) and quantification (d) of cable recoil following laser ablation in control and gCre;Fl/Fl embryos. Points represent individual embryos. Scale bar = 5 µm. **e.** 3D volumetric rendering of reflection images showing pseudo-coloured protrusions at or adjacent to the zippering point in control and gCre;Fl/Fl embryos. **f.** Quantification of filopodial-type protrusion density in control and gCre;Fl/Fl embryos. Points represent individual embryos. **g-h.** Surface subtracted (showing only the SE) high-resolution images of adherens junctions showing equivalent non-phosphorylated β-catenin co-localising with F-actin at cell borders (i) and tight junctions showing ZO-1 (j) in control and gCre;Fl/Fl embryos. **h.** Arrowheads show cell-cell borders rostral to the zippering point, circles indicate enrichment at tri-cellular junctions in both genotypes. Scale bars = 10 µm. **i.** Quantification of border constriction ratios at the indicated cell position in control and gCre;Fl/Fl embryos with 17-20 somites. Points represent individual embryos, averaging their left and right cells in the semi-rosette. The illustration schematizes the pattern of border constriction, with diminished constriction in position 1 of gCre;Fl/Fl embryos (indicated by green arrows). Two-way ANOVA (genotype, position) with post-hoc T-tests within genotype.

Zippering point protrusions may form cell-cell or cell-ECM attachments. The ECM substrate fibronectin is present in both gCre;Fl/Fl and control embryos (Figure S3c). New cell-cell junctions form between cells which meet at the midline (Figure 5a’’) and both adherens and tight junction components are identifiable along these midline cell borders (Figure 5g-h). Cells engaged in the supracellular cables progressively constrict their leading-edge border as they approach the zipper (numbered in sequential rows with those in position 1 being at the zippering point, Figure 5a’’’). Both control and gCre;Fl/Fl embryos show significant border constriction between cells in position 3 versus position 2 (Figure 5i). However, whereas position 1 cells of control embryos constrict further, in gCre;Fl/Fl embryos they do not, and are significantly less constricted than in controls (Figure 5i). Collectively, these analyses exclude various potential causes of slow zippering, including diminished cable constriction, presence of midline junctions, fibronectin substrate, or presence of cellular protrusions. The two abnormalities observed in gCre;Fl/Fl embryos are over-abundance of filopodial protrusions and diminished border constriction at the zippering point.

### Loss of CFL1 diminishes filopodial dynamicity

High-resolution imaging of the zippering point does not identify cytoskeletal or cell-cell junction differences between control and gCre:Fl/Fl embryos which might account for diminished leading-edge constriction. Both assemble continuous cables bordering F-actin-rich protrusions (Figure 5b, 6a). Both have interspersed adherens junctions punctuating the cables (Figure 6b). We therefore sought to assess dynamic behaviours of filopodial protrusions to test whether their over-abundance in gCre;Fl/Fl embryos reflects their function. To achieve this, we first used high-resolution live imaging of the zippering point to quantitatively assess protrusion dynamics in wildtype embryos (Figure 6c). We observe dynamic exploratory behaviours of protrusions which appear to make transient contacts across the midline then regress, whereas other ‘pioneer’ protrusions form contacts which are then stabilised (Figure S4a). Both lamellipodial (Figure S4b) and filopodial (Figure 6c, Figure S4c) protrusions are observed forming pioneer contacts. However, we also observe substantial differences in the persistence of the two types of protrusions: whereas lamellipodial protrusions are stable features (Figure 6d), most filopodial ones are only observed for up to two minutes in wildtype embryos (Figure 6e).

**Figure 6:**
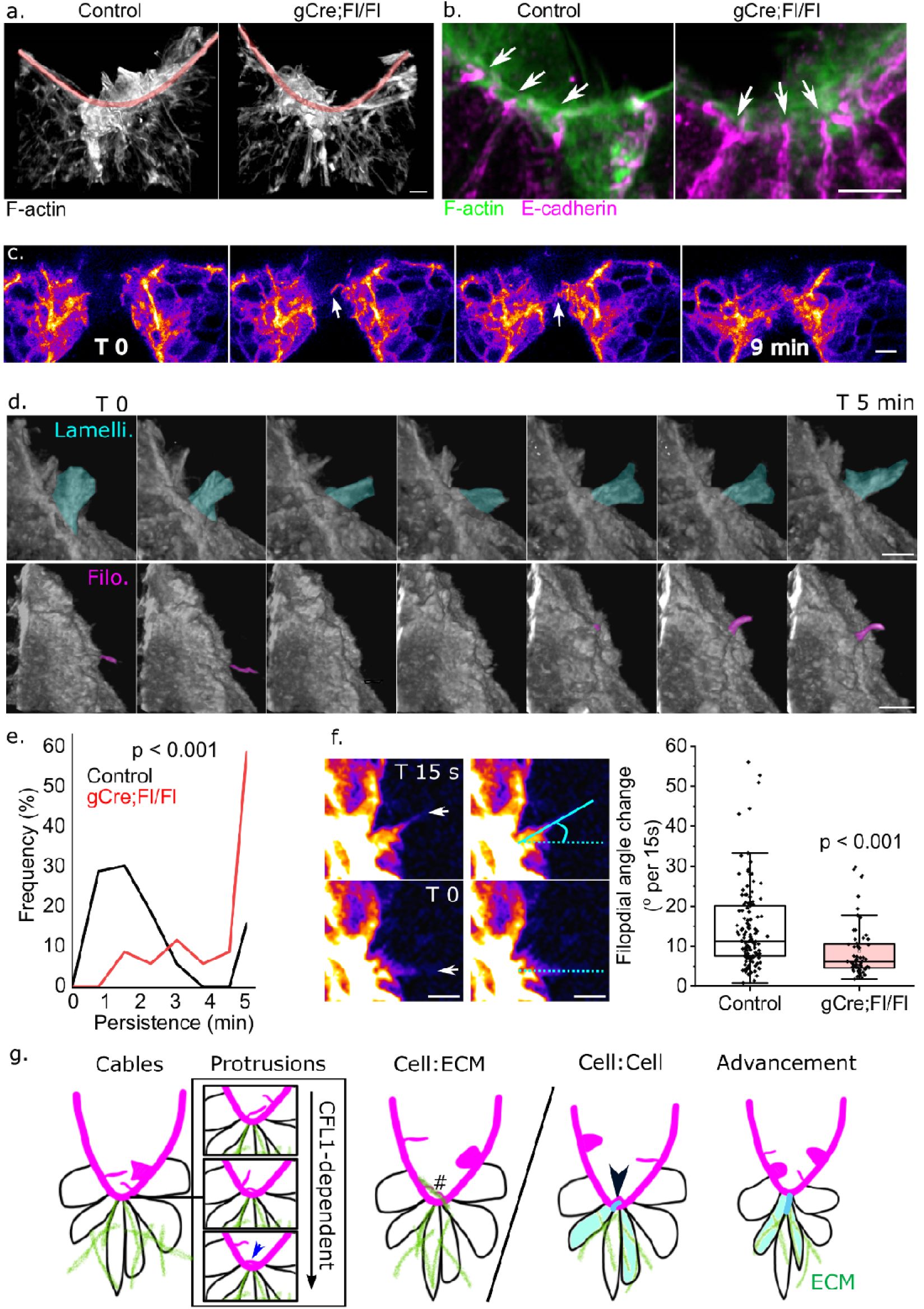
Assessment of cellular protrusion dynamics during zippering. **a.** 3D reconstruction of AiryScan super-resolution images of the zippering point in control and gCre;Fl/Fl embryos showing equivalent abundance of protrusions directly at this point, overlapping with the supracellular cables indicated by the red highlight. Scale bar = 5 µm. **b.** AiryScan visualisation of E-cadherin-labelled adherens junctions interspersed into the supracellular cable equivalents in control and gCre;Fl/Fl embryos. Scale bar = 5 µm. **c.** Live imaging of protrusion dynamics directly at the zippering point, showing a filopodial type pioneer protrusion (arrow) which forms an initial contact which is subsequently stabilised by broader cell outgrowth. Images are processed with PureDenoise and shown in Fire LUT to aid visualisation of delicate structures. Scale bar = 10 µm. **d.** 3D pseudo-coloured live imaging illustrating dynamics of lamellipodial and filopodial protrusions adjacent to the zippering point. Scale bar = 10 µm. **e.** Frequency plot quantifying the persistence of filopodial protrusions in control and gCre;Fl/Fl embryos (155 filopodia from 13 control embryos and 63 filopodia from 7 gCre;Fl/Fl embryos). **f.** Visualisation and quantification of the change in angle (cyan lines) of filopodial protrusions (white arrow) between live-imaged snapshots of a control embryo. Points represent the average change in angle of each protrusion over the time course it was visualised for (128 filopodia from 11 control embryos and 62 filopodia from 7 gCre;Fl/Fl embryos). Scale bar = 5 µm. **g.** Schematic representation of the proposed spinal zippering sequence. SE cells assemble contractile cables and extend protrusions. These protrusions undergo dynamic displacements until ones close to the zippering point establish new contacts across the embryonic midline. These pioneer protrusions can form cell-cell or cell-ECM contacts which are then stabilised. The midline cells then constrict their leading-edge border as they leave the zippering point.

Control and gCre;Fl/Fl embryos were compared using short-term, high-frequency and high-resolution imaging necessary to capture zippering point Z-stacks and analyse filopodial movement of multiple embryos per litter. Filopodia in gCre;Fl/Fl embryos are significantly more stable, often persisting longer than the 5-minute imaging periods selected based on wild-type data (Figure 6e). However, they are also significantly less dynamic, showing smaller changes in angle between timepoints (Figure 6f). Thus, our live imaging analyses provides evidence that filopodial exploratory displacements precede establishment of pioneer contacts across the embryo midline, and that this exploratory function is diminished without CFL1 (Figure 6g).

## Discussion

Failure of epithelial fusion causes clinically important congenital malformations including coloboma, cleft lip/palate and neural tube defects^1^. Yet, the sequence of cell behaviours by which abutting epithelia undergo fusion are poorly delineated, particularly in mammals. We propose a sequence of five steps based on their caudal-to-rostral position (Figure 6g) and present their qualitative or quantitative assessment during mouse spinal zippering-mediated fusion: *i.* cable constriction, *ii.* protrusion extension and exploratory movements forming pioneer contacts, *iii.* cell-cell and *iv.* cell-ECM adhesion remodelling, followed by *v.* zipper advancement through shrinking of the leading-edge cell borders. It is likely that additional steps or inter-relations between identified behaviours will be added in future work, and that some will not be applicable to other fusion events. For example, chicken embryos close their spinal neural tube without assembling supracellular cables at the edge of their SE^56^.

We apply this paradigm to specifically test the roles of CFL1 in zippering. Global deletion of *Cfl1* has previously been found to disrupt apical enrichment of F-actin in both the cranial and spinal neuroepithelium^38,39^ and results in up-regulation of destrin^40^ which might compensate for some of its functions. Global *Cfl1^-/-^* embryos are similar to wildtype controls at embryonic day 9, but die over the next two days^40^. Surprisingly, we find that deletion of *Cfl1* in the neuroepithelium throughout the period of spinal closure did not impair progression of closure. The mild phenotype in our *Cdx2^Cre^* deletion model suggests CFL1 is likely to play critical roles in tissues unrecombined in this model, such as the heart^14^. Our *Grhl3^Cre^*model also avoids severe stunting during neural tube closure, making it mechanistically tractable, unlike deletion of *Cdc42* with the same Cre-driver^10^. Although we observe larger aggregates of apoptotic bodies in the dorsal midline of *Grhl3^Cre/+^Cfl1^Fl/Fl^*embryos, wildtype embryos also have abundant dorsal midline apoptosis^19^ and aggregates are not observed adjacent to the zippering point in either genotype. We therefore consider them unlikely to impact zippering. Apoptotic aggregates may be bigger in conditional knockouts because of changes in CFL1-dependant formation of blebs by the apoptotic cells themselves^57^, or diminished F-actin remodelling in the apoptotic and surrounding cells which extrude them^58^.

F-actin dysregulation, producing more prominent stress fibres in SE cells lacking CFL1, is consistent with its known roles in cultured cells^54^. The moderate yet highly significant negative corelation between pCFL1 and F-actin intensity we observe in SE cells of wildtype embryos was unexpected. Stable interventions which decrease pCFL1 also decrease F-actin in cultured cells^59^ and loss of its phosphatase increases both pCFL1 and F-actin intensity in *Drosophila*^60^ (positive correlations). Possible explanations for this discrepancy include differences in pCFL1/F-actin dynamics inferred from a tissue snapshot. Paradoxically, at high cofilin:actin ratios, CFL1 can promote F-actin assembly^61^, such that cells with high pCFL1 could be those with sufficiently low CFL1 activity to sever F-actin. These alternatives are difficult to disentangle *in vivo* with the tools available, and we leave them for future work.

We provide new insights into *in vivo* cell- and tissue-level regulation of CFL1 activity through widespread ROCK-dependent phosphorylation superimposed on sub-cellular localisation and cell cycle- or tissue region-specific phosphorylation (see Figure S5 for hypothesised molecular mechanisms regulating zippering). Particularly striking is the enrichment of total CFL at the leading edge of SE cells adjacent to the zippering point, but only at earlier somite stages corresponding to when CDC42 activity promotes filopodial extension^10^. The molecular bases for CFL1 sub-cellular localisation and Rho-GTPAse switching during spinal zippering is unknown and we intend to explore them in future work. Previous *in* vitro studies observed recruitment of CFL1 to the leading edge of cells extending filopodial protrusions, followed by localisation of CFL1 into filopodia as they retract^62^. An increase in filopodia has also been reported in cultured CFL1-depleted mesenchymal cells^55^. Together with our observation of excessive number and persistence of filopodia in embryos lacking CFL1, including over the closed portion of the neural tube in some embryos, we conclude that CFL1 normally destabilises these structures during spinal zippering. Filopodial disassembly may be an important step in the transition from CDC42-driven partially-mesenchymal protrusive activity at the neuropore margin, to fully epithelial SE character producing Rho-dependent CFL1 phosphorylation after leaving the zipper. Both suppression and accentuation of SE epithelial properties, as indicated by their levels of cell-cell adhesion proteins, can stall spinal zippering^11^.

CFL1-dependent assembly of a peri-junctional F-actin belt facilitates formation of new cell-cell adhesions necessary to diminish epithelial permeability in gut epithelial cells^63^. Both adherens and tight junctions are present between midline SE cells which recently left the zippering point in *Cfl1* conditional knockouts, suggesting CFL1 is not required for their assembly. Interpretation of this finding is tempered by the slower rate of zippering in these embryos, such that nascent cell-cell (and cell-ECM^64^) junctions may have more time to mature. Direct analysis of cell-ECM adhesion is limited, but we consider it unlikely to explain failure of zippering because equivalent deletion of the integrin *Itgb1* impairs closure at later somite stages (>22 somites)^19^ than in our *Cfl1-*deletion model. SE CFL1 is lost by those later stages in our *Cdx2^Cre^* model, but closure is not impaired.

CFL1-mediated F-actin turnover is required to connect the cytoskeleton to adherens junctions and transmit tension between cells in an apically-constricting *Drosophila* epithelium^65^. In contrast, we do not observe quantifiable reductions in junctional or cable tension in SE cells lacking CFL1. High tension structures such as the supracellular cables may be shielded from the effects of CFL1 because F-actin filaments under tension are protected from being severed^66^. Consistent with having equivalent junctional tension, we do not observe changes in YAP nuclear localisation as a molecular readout of SE mechanics^24,55^. This contrasts with *in vitro* findings that CFL1 knockdown increases YAP nuclear localisation in confluent mesenchymal cells^55^. Some of these discrepancies may be explained by *in vitro* versus *in vivo* comparisons (e.g. stiff versus soft substrates) or species differences (e.g. tight junctions present in mouse but not in *Drosophila*^67^). Evolutionary conservation is also apparent. Zippering during *Drosophila* dorsal closure bears several similarities to mouse spinal zippering, including formation of leading edge cables and extension of protrusions which contact genetically-patterned partners, interdigitate, and form new cell-cell junctions^68^. SE protrusions have also been live-imaged during mouse cranial neural tube closure, where they appear to form long ‘bridges’^69^. These clearly differ from the dynamic, transient or pioneer filopodial protrusions we live-image in the spine.

Pioneer protrusions might initiate bi-directional signalling through ephrin/Eph receptors^70^, form cell-ECM contacts, establish nascent cell-cell adhesions, and/or provide a medial pulling force^68^. A SE-specific tight junction protein has recently been shown to facilitate progression of spinal buttoning in chicken embryos by dampening SE tension^12^, and some components of tight junctions are identifiable on mouse zippering protrusions suggesting they are poised to form new adhesions^11^. In contrast, we do not observe adherens junction proteins in protrusions, and E-cadherin neutralisation in cultured embryos does not increase PNP length (a hallmark of slower zippering)^24^. In other systems, filopodial protrusions are stabilised by matching homotypic adhesion molecules on opposing cells^5^, or by actively indenting the membrane of a cell without matching filopodia^71^. Other species close their neural tubes without equivalent structures, including chicken embryos in which sparse bleb-like protrusions are the predominant type observed during spinal buttoning^7^.

Our findings contrast with an alternative model of mouse spinal zippering which proposes that protrusion convergence generates semi-rosette cell arrangements and the force needed to pull the two halves toward the midline^20^. We do not observe ‘pinching in’ of the neural folds adjacent to protrusions are predicted by that model, and the junctional reduction of cells in position 2 but not position 1 of our conditional *Cfl1* knockout is also inconsistent with that model. The junctional constriction pattern we observe here is also distinct from the effect of *Itgb1* deletion; in which junctional constriction is diminished at multiple cell positions caudal to the zippering point, but only at later stages of development^19^. Both the *Itgb1* and current studies suggest that formation/stabilisation of adhesions precedes semi-rosette junctional constriction, but we cannot conclude from our imaging whether pioneering contacts trigger junctional shortening of cells at the zippering point. This would be consistent with the equivalent process in *Ciona*, in which junctional constriction is triggered by remodelling of adherens junctions^17^. However, whereas in *Ciona* only the cells at ‘position 1’ constrict their leading-edge border^17,18^, in mice there is constriction between position 3 to position 2, then between position 2 and 1 in control embryos^19^. The 3-to-2 constriction is independent of CFL1, whereas only wildtype embryos show significant 2-to-1 constriction.

In conclusion, we propose a spatially-organised sequence of cell behaviours necessary for progression of mammalian spinal zippering (Figure 6g). Applying this framework, we identify novel functions for CFL1 in epithelial fusion by enhancing filopodial dynamicity and maintaining constriction of midline SE cell borders at the zippering point. Aggregation of these effects – of which we infer diminished filopodial dynamicity is the likely inciting defect – slows zippering speed by ∼30%, without halting it entirely in any embryos tested. The consequence is a reduced likelihood of completing spinal neural tube closure, leading to partially penetrant spina bifida.

## Materials And Methods

### Mouse transgenic alleles

All mouse lines were generated or maintained on a C57/BL6 background. Genotyping was performed by PCR on DNA extracted from ear clips. The following transgenic mouse lines were used: *Cdx2^Cre^* ^72^ (gene symbol: Tg (CDX2-cre)101Erf, MGI: 3696953) and *Grhl3^Cre^* ^73^ (gene symbol: Grhl3^tm1^ (cre)^Cgh^, MGI: 4430902).

Mice with a conditional floxed allele of *Cfl1* (*Cfl1^Fl/Fl^*) were derived from the strain C57BL/6N-Atm1Brd Cfl1tm1a (KOMP)Mbp/JMmucd (RRID:MMRRC_047079-UCD), obtained from the Mutant Mouse Resource and Research Center (MMRRC) at University of California at Davis, an NIH-funded strain repository, and was donated to the MMRRC by The KOMP Repository, UC Davis Mouse Biology Program (Stephen Murray, Ph.D., The Jackson Laboratory). Mice with the tm1a allele were crossed with flipase-expressing mice to generate the conditional tm1c allele. The flipase allele used was derived from C57BL/6N-Tg (CAG-Flpo)1Afst/Mmucd (RRID:MMRRC_036512-UCD) obtained from the MMRRC, and was donated by the MMRRC at UC Davis. The original transgenic was donated by Dr. Konstantinos Anastassiadis from Technische Universitaet Dresden.

Homozygous *Cfl1^fl/fl^* mice were crossed with Cre-heterozygous animals, to generate *Cfl1^fl/+^*;*Cre/+* animals. These were in turn crossed with *Cfl1^fl/fl^* mice to generate experimental *Cfl1^fl/fl^*;*Cre/+* embryos.

### Animal procedures

All animal work was performed under the regulation of the UK Animals (Scientific Procedures) Act 1986 and the National Centre for the 3Rs’ Responsibility in the Use of Animals for Medical Research (2019). Mice used for breeding and experimental mating were no less than 8 weeks and no more than 1 year of age.

Mice were mated overnight or early in the morning, and checked for a copulation plug the next morning or at midday respectively. Positive plugs found after overnight breeding were designated embryonic day (E) 0.5, while plugs found the same day were considered E0.5 from midnight. Recombination domains were visualised by crossing heterozygous Cre mice with homozygous *Rosa26-mTmG* mice.

Pregnant dams were sacrificed by cervical dislocation at appropriate embryonic stages, from E8.5 – 14.5. Individual deciduas were dissected from the uterus in warm Dulbecco’s Modified Eagle’s Medium containing 25 mM HEPES and 10% fetal calf serum. Embryos were then dissected free of decidua, and used for culture, fixation or laser ablation experiments.

### Embryo culture

Embryo culture was performed as previously described ^74^. Decidua, trophoblast and Reichert’s membrane layers were carefully removed, excluding the trophoblast of the ectoplacental cone and taking care not to pierce the yolk sac. Embryos were then stage matched and randomly allocated to experimental groups. Embryos were carefully added to a 30 ml culture tube with pre-warmed rat serum filtered through a 0.45 µm Millipore filter (0.5 ml per embryo). Culture tubes were gassed for 1 min with 20% O_2_, 5% CO_2_, 75% N_2_, sealed using vacuum grease and added to a roller culture incubator for the duration of the experiment. After cultures, embryos were processed as below for fixing. RHO/ROCK inhibitor (Y27632, Cambridge Biosciences SM02-1) dissolved in water was added to rat serum at a concentration of 10 µM.

To measure the speed of zippering point progression in culture as previously described^11^, a small hole was created in the yolk sac and amnion using dissection forceps, revealing the zippering point. A small volume of DiI (Invitrogen V22889) was injected lateral to the zippering point, and embryos were added to rat serum and cultured for 6.5 h. Zippering speed was calculated from the distance moved by the zippering point beyond the DiI mark, divided by elapsed time.

Embryos were fixed by removing all extra-embryonic layers including the yolk sac and amnion washing briefly in ice cold PBS and fixed immediately in 4% PFA in PBS on ice. Yolk sacs were also washed in PBS and kept for genotyping. Yolk sacs (embryos) or ear clips (mice) were lysed for at least 4 h at 56°C in 50 µl lysis buffer (Viagen Biotech) with 10% proteinase K. Lysates were diluted 1:10 with Milli-Q water and processed for PCR using GoTaq G2 Flexi DNA Polymerase kit (Promega). Reactions were placed in a thermocycler on a touchdown program, and subsequent amplified DNA was evaluated by gel electrophoresis.

### Immunofluorescence

Primary and secondary antibodies used in this study are detailed in Supplementary Table 1. All washes and incubation steps were on a benchtop shaker. For ROCK1 and pMLC-II primary antibodies, an antigen retrieval step was included: embryos were preincubated for 15 min in sodium citrate solution (10 mM in PBS + 0.05% Tween 20, pH 6) at room temperature (RT), followed by 25 min at 90°C in fresh sodium citrate solution. Embryos were allowed to cool at RT before proceeding to the blocking step. For all other antibodies, embryos were permeabilised in PBS + 0.1% Triton (PBST) for 1 h at RT.

After antigen retrieval or permeabilisation, embryos were incubated in blocking solution (PBST + 5% bovine serum albumin), either overnight at 4°C or for 6 h at RT. Primary antibodies were diluted in blocking solution and incubated at 4°C overnight. Primary antibodies were diluted at the following dilutions: 1;100 rabbit anti-CFL1, 1:200 rabbit anti-pCFL1, 1:100 rabbit anti-ROCK1, 1:200 mouse anti-pHH3, 1:200 rabbit anti-pMLC-II, mouse anti-E cadherin, 1:200 rabbit anti-cCasp3. The following day, primary antibody solutions were removed and embryos were washed 3×1 h in blocking solution at RT. Embryos were then incubated in secondary antibody solution for 2 h at RT. When used, DAPI and/or Phalloidin were also added during this step. Secondary antibodies were diluted in blocking solution at the following concentrations: Alexa fluor -405 and -647 secondary antibodies, 1:250; Alex fluor -488 and -568, 1:500. DAPI was diluted at 1:5000, and Phalloidin at 1:250. Finally, embryos were washed 1x 1 h in blocking solution, 1x 1 h in PBST, 1 h in PBS, and stored in PBS + 1% azide.

### Confocal microscopy and laser ablation

All confocal imaging was performed using a Zeiss Examiner LSM880 confocal microscope using 10x/NA0.5/ or 20x/NA1.0 Plan Apochromat water immersion objectives. Laser power and gain settings were kept the same for embryos from a single experiment.

For laser ablation, embryos were stained, positioned and ablated as previously described^75^. After dissection, yolk sacs were kept for genotyping and live embryos were stained for 5 min in 1:500 Cell Mask (Thermo Fisher Scientific) in DMEM (without FBS) at 37 C. The caudal half of the embryo was separated using dissection forceps, as movement caused by the heartbeat prevents accurate ablations, and positioned in a pre-warmed agarose plate with DMEM. Curved microsurgical needles were positioned in the agarose to provide support to the tissue and keep it stable during imaging. The tissue was positioned with the dorsal side facing up; the PNP or caudal surface ectoderm was positioned as flat as possible, depending on whether the cable or surface ectoderm cells were being ablated. Cell borders were imaged in a single z plane. After 1 sec imaging, a single cell border was ablated using a MaiTai laser (710 nm wavelength, 80% laser power, 0.34 µsec pixel dwell time, 20 iterations).

### Image analysis

All image processing and analysis was carried out using FIJI^76^. Images which are ‘surface subtracted’^16^ to selectively visualize the surface ectoderm have a homogenous, null background where there is no surface. 3D projections are generated in FIJI as 8-bit images and also have a homogenous, computer background. Brightness and contrast were adjusted evenly across all image panes. Salt-and-pepper noise was eliminated using the remove function in FIJI when appropriate. AiryScan images underwent standard processing in ZEN Blue using the manufacturer’s software. The ‘surface subtraction’ macro used for extracting surface signal from fluorescence images is available on Github courtesy of Dr Dale Moulding https://github.com/DaleMoulding/Fiji-Macros.

Fluorescence intensity quantifications of an area, intensity profiles, length and area measurements were made using standard tools in FIJI. Initial recoil of laser ablations was calculated as the change in length of a line drawn between two reference points before and after the ablation^75^. Protrusion analysis was carried out using raw z stacks. Individual protrusions were identified and recorded using the ROI manager tool. Protrusions were then tracked over time - protrusions could move in Z as well as X and Y, so they were also tracked over z slices. Image clarity was computationally improved using PureDenoise when appropriate.

### Statistical analysis

Statistical analysis was performed using OriginLab and Microsoft Excel software. Two-tailed p ≤ 0.05 was considered statistically significant. The embryo was the unit of measure, except for live imaging analysis in which individual protrusions were the unit of measure. Slopes were compared by F-test, and means of two groups by Student’s T-test or Mann Whitney U test. Comparison of multiple groups was by ANOVA with post-hoc Bonferroni. Where possible, analyses were performed blind to genotyping or treatment group.

## Acknowledgements

This study was supported by the Wellcome Trust (211112/Z/18/Z and 211112/Z/18/A to G.L.G.), GOSH CC (VS2505 to G.L.G) and the Royal Society (RG\R2\232082 to GLG). E.M. was funded by a Marie Skłodowska-Curie Actions project (101067028).

## Author contributions

Conception or design of the work: GLG. Acquisition, analysis, or interpretation: GLG, ARM, AK, HC, RS, EM, NDEG, AJC. Drafting the work: GLG, ARM. Reviewing: GLG, AK, HC, RS, EM, NDEG, AJC. Final approval and agreement to be accountable: All authors.

## References

1 Chan, B. H. C., Moosajee, M. & Rainger, J. Closing the Gap: Mechanisms of Epithelial Fusion During Optic Fissure Closure. Front Cell Dev Biol 8, 620774 (2020). 10.3389/fcell.2020.620774

2 Pasakarnis, L., Frei, E., Caussinus, E., Affolter, M. & Brunner, D. Amnioserosa cell constriction but not epidermal actin cable tension autonomously drives dorsal closure. Nat Cell Biol 18, 1161–1172 (2016). 10.1038/ncb3420

3 Ducuing, A. & Vincent, S. The actin cable is dispensable in directing dorsal closure dynamics but neutralizes mechanical stress to prevent scarring in the Drosophila embryo. Nat Cell Biol 18, 1149–1160 (2016). 10.1038/ncb3421

4 Swope, D., Kramer, J., King, T. R., Cheng, Y. S. & Kramer, S. G. Cdc42 is required in a genetically distinct subset of cardiac cells during Drosophila dorsal vessel closure. Dev Biol 392, 221–232 (2014). 10.1016/j.ydbio.2014.05.024

5 Zhang, S., Teng, X., Toyama, Y. & Saunders, T. E. Periodic Oscillations of Myosin-II Mechanically Proofread Cell-Cell Connections to Ensure Robust Formation of the Cardiac Vessel. Curr Biol 30, 3364–3377 e3364 (2020). 10.1016/j.cub.2020.06.041

6 King, T. R., Kramer, J., Cheng, Y. S., Swope, D. & Kramer, S. G. Enabled/VASP is required to mediate proper sealing of opposing cardioblasts during Drosophila dorsal vessel formation. Dev Dyn 250, 1173–1190 (2021). 10.1002/dvdy.317

7 Legere, E. A. et al. Claudin-3 in the non-neural ectoderm is essential for neural fold fusion in chicken embryos. Dev Biol 507, 20–33 (2024). 10.1016/j.ydbio.2023.12.009

8 Van Straaten, H. W., Janssen, H. C., Peeters, M. C., Copp, A. J. & Hekking, J. W. Neural tube closure in the chick embryo is multiphasic. Dev Dyn 207, 309–318 (1996). 10.1002/(SICI)1097-0177(199611)207:3<309::AID-AJA8>3.0.CO;2-L

9 Copp, A. J. Relationship between timing of posterior neuropore closure and development of spinal neural tube defects in mutant (curly tail) and normal mouse embryos in culture. J Embryol Exp Morphol 88, 39–54 (1985).

10 Rolo, A. et al. Regulation of cell protrusions by small GTPases during fusion of the neural folds. Elife 5, e13273 (2016). 10.7554/eLife.13273

11 Nikolopoulou, E. et al. Spinal neural tube closure depends on regulation of surface ectoderm identity and biomechanics by Grhl2. Nat Commun 10, 2487 (2019). 10.1038/s41467-019-10164-6

12 Legere, E.-A., Dumont, M., Yamanaka, Y., Galea, G. L. & Ryan, A. K. Increased tissue tension caused by depletion of CLDN3 in the non-neural ectoderm causes neural fold fusion defects in chick embryos. bioRxiv, 2025.2010.2014.682463 (2025). 10.1101/2025.10.14.682463

13 Santos, C. et al. Spinal neural tube formation and tail development in human embryos. Elife 12 (2024). 10.7554/eLife.88584

14 Maniou, E. et al. Caudal Fgfr1 disruption produces localised spinal mis-patterning and a terminal myelocystocele-like phenotype in mice. Development 150 (2023). 10.1242/dev.202139

15 Epstein, D. J., Vekemans, M. & Gros, P. Splotch (Sp2H), a mutation affecting development of the mouse neural tube, shows a deletion within the paired homeodomain of Pax-3. Cell 67, 767–774 (1991). 10.1016/0092-8674(91)90071-6

16 Galea, G. L. et al. Vangl2 disruption alters the biomechanics of late spinal neurulation leading to spina bifida in mouse embryos. Dis Model Mech 11 (2018). 10.1242/dmm.032219

17 Hashimoto, H. & Munro, E. Differential Expression of a Classic Cadherin Directs Tissue-Level Contractile Asymmetry during Neural Tube Closure. Dev Cell 51, 158–172 e154 (2019). 10.1016/j.devcel.2019.10.001

18 Hashimoto, H., Robin, F. B., Sherrard, K. M. & Munro, E. M. Sequential contraction and exchange of apical junctions drives zippering and neural tube closure in a simple chordate. Dev Cell 32, 241–255 (2015). 10.1016/j.devcel.2014.12.017

19 Mole, M. A. et al. Integrin-Mediated Focal Anchorage Drives Epithelial Zippering during Mouse Neural Tube Closure. Dev Cell 52, 321–334 e326 (2020). 10.1016/j.devcel.2020.01.012

20 Zhou, C. J. et al. Non-neural surface ectodermal rosette formation and F-actin dynamics drive mammalian neural tube closure. Biochem Biophys Res Commun 526, 647–653 (2020). 10.1016/j.bbrc.2020.03.138

21 Galea, G. L. et al. Biomechanical coupling facilitates spinal neural tube closure in mouse embryos. Proc Natl Acad Sci U S A 114, E5177–E5186 (2017). 10.1073/pnas.1700934114

22 Maniou, E. et al. Hindbrain neuropore tissue geometry determines asymmetric cell-mediated closure dynamics in mouse embryos. Proc Natl Acad Sci U S A 118 (2021). 10.1073/pnas.2023163118

23 Butler, M. B. et al. Rho kinase-dependent apical constriction counteracts M-phase apical expansion to enable mouse neural tube closure. J Cell Sci 132 (2019). 10.1242/jcs.230300

24 Marshall, A. R., Galea, G. L., Copp, A. J. & Greene, N. D. E. The surface ectoderm exhibits spatially heterogenous tension that correlates with YAP localisation during spinal neural tube closure in mouse embryos. Cells Dev 174, 203840 (2023). 10.1016/j.cdev.2023.203840

25 Jaffe, E. & Niswander, L. Loss of Grhl3 is correlated with altered cellular protrusions in the non-neural ectoderm during neural tube closure. Dev Dyn 250, 732–744 (2021). 10.1002/dvdy.292

26 Van Aelst, L. & Symons, M. Role of Rho family GTPases in epithelial morphogenesis. Genes Dev 16, 1032–1054 (2002). 10.1101/gad.978802

27 Chauhan, B. K., Lou, M., Zheng, Y. & Lang, R. A. Balanced Rac1 and RhoA activities regulate cell shape and drive invagination morphogenesis in epithelia. Proc Natl Acad Sci U S A 108, 18289–18294 (2011). 10.1073/pnas.1108993108

28 Machesky, L. M. & Hall, A. Role of actin polymerization and adhesion to extracellular matrix in Rac- and Rho-induced cytoskeletal reorganization. J Cell Biol 138, 913–926 (1997). 10.1083/jcb.138.4.913

29 Nobes, C. D. & Hall, A. Rho, rac, and cdc42 GTPases regulate the assembly of multimolecular focal complexes associated with actin stress fibers, lamellipodia, and filopodia. Cell 81, 53–62 (1995). 10.1016/0092-8674(95)90370-4

30 Chen, T. J., Gehler, S., Shaw, A. E., Bamburg, J. R. & Letourneau, P. C. Cdc42 participates in the regulation of ADF/cofilin and retinal growth cone filopodia by brain derived neurotrophic factor. J Neurobiol 66, 103–114 (2006). 10.1002/neu.20204

31 Kanellos, G. & Frame, M. C. Cellular functions of the ADF/cofilin family at a glance. J Cell Sci 129, 3211–3218 (2016). 10.1242/jcs.187849

32 Pyronneau, A. et al. Aberrant Rac1-cofilin signaling mediates defects in dendritic spines, synaptic function, and sensory perception in fragile X syndrome. Sci Signal 10 (2017). 10.1126/scisignal.aan0852

33 Soosairajah, J. et al. Interplay between components of a novel LIM kinase-slingshot phosphatase complex regulates cofilin. EMBO J 24, 473–486 (2005). 10.1038/sj.emboj.7600543

34 Edwards, D. C., Sanders, L. C., Bokoch, G. M. & Gill, G. N. Activation of LIM-kinase by Pak1 couples Rac/Cdc42 GTPase signalling to actin cytoskeletal dynamics. Nat Cell Biol 1, 253–259 (1999). 10.1038/12963

35 Agarwal, P. & Zaidel-Bar, R. Principles of Actomyosin Regulation In Vivo. Trends Cell Biol 29, 150–163 (2019). 10.1016/j.tcb.2018.09.006

36 Koenderink, G. H. & Paluch, E. K. Architecture shapes contractility in actomyosin networks. Curr Opin Cell Biol 50, 79–85 (2018). 10.1016/j.ceb.2018.01.015

37 Ikeda, S. et al. Aberrant actin cytoskeleton leads to accelerated proliferation of corneal epithelial cells in mice deficient for destrin (actin depolymerizing factor). Hum Mol Genet 12, 1029–1037 (2003). 10.1093/hmg/ddg112

38 Escuin, S. et al. Rho-kinase-dependent actin turnover and actomyosin disassembly are necessary for mouse spinal neural tube closure. J Cell Sci 128, 2468–2481 (2015). 10.1242/jcs.164574

39 Grego-Bessa, J., Hildebrand, J. & Anderson, K. V. Morphogenesis of the mouse neural plate depends on distinct roles of cofilin 1 in apical and basal epithelial domains. Development 142, 1305–1314 (2015). 10.1242/dev.115493

40 Gurniak, C. B., Perlas, E. & Witke, W. The actin depolymerizing factor n-cofilin is essential for neural tube morphogenesis and neural crest cell migration. Dev Biol 278, 231–241 (2005). 10.1016/j.ydbio.2004.11.010

41 Bamburg, J. R., Harris, H. E. & Weeds, A. G. Partial purification and characterization of an actin depolymerizing factor from brain. FEBS Lett 121, 178–182 (1980). 10.1016/0014-5793(80)81292-0

42 Goley, E. D., Rodenbusch, S. E., Martin, A. C. & Welch, M. D. Critical conformational changes in the Arp2/3 complex are induced by nucleotide and nucleation promoting factor. Mol Cell 16, 269–279 (2004). 10.1016/j.molcel.2004.09.018

43 Bergert, M., Chandradoss, S. D., Desai, R. A. & Paluch, E. Cell mechanics control rapid transitions between blebs and lamellipodia during migration. Proc Natl Acad Sci U S A 109, 14434–14439 (2012). 10.1073/pnas.1207968109

44 Lappalainen, P., Kotila, T., Jegou, A. & Romet-Lemonne, G. Biochemical and mechanical regulation of actin dynamics. Nat Rev Mol Cell Biol 23, 836–852 (2022). 10.1038/s41580-022-00508-4

45 Buracco, S., Claydon, S. & Insall, R. Control of actin dynamics during cell motility. F1000Res 8 (2019). 10.12688/f1000research.18669.1

46 Fort, L. et al. Fam49/CYRI interacts with Rac1 and locally suppresses protrusions. Nat Cell Biol 20, 1159–1171 (2018). 10.1038/s41556-018-0198-9

47 Hu, K., Ji, L., Applegate, K. T., Danuser, G. & Waterman-Storer, C. M. Differential transmission of actin motion within focal adhesions. Science 315, 111–115 (2007). 10.1126/science.1135085

48 Redd, M. J., Cooper, L., Wood, W., Stramer, B. & Martin, P. Wound healing and inflammation: embryos reveal the way to perfect repair. Philos Trans R Soc Lond B Biol Sci 359, 777–784 (2004). 10.1098/rstb.2004.1466

49 Wang, A., Ma, X., Conti, M. A. & Adelstein, R. S. Distinct and redundant roles of the non-muscle myosin II isoforms and functional domains. Biochem Soc Trans 39, 1131–1135 (2011). 10.1042/BST0391131

50 Elam, W. A., Kang, H. & De la Cruz, E. M. Biophysics of actin filament severing by cofilin. FEBS Lett 587, 1215–1219 (2013). 10.1016/j.febslet.2013.01.062

51 Oosterheert, W., Boiero Sanders, M., Hofnagel, O., Bieling, P. & Raunser, S. Choreography of rapid actin filament disassembly by coronin, cofilin, and AIP1. Cell 188, 6845–6860 e6827 (2025). 10.1016/j.cell.2025.09.016

52 Yonezawa, N., Nishida, E., Iida, K., Yahara, I. & Sakai, H. Inhibition of the interactions of cofilin, destrin, and deoxyribonuclease I with actin by phosphoinositides. J Biol Chem 265, 8382–8386 (1990).

53 Zhao, H., Hakala, M. & Lappalainen, P. ADF/cofilin binds phosphoinositides in a multivalent manner to act as a PIP(2)-density sensor. Biophys J 98, 2327–2336 (2010). 10.1016/j.bpj.2010.01.046

54 Kanellos, G. et al. ADF and Cofilin1 Control Actin Stress Fibers, Nuclear Integrity, and Cell Survival. Cell Rep 13, 1949–1964 (2015). 10.1016/j.celrep.2015.10.056

55 Aragona, M. et al. A mechanical checkpoint controls multicellular growth through YAP/TAZ regulation by actin-processing factors. Cell 154, 1047–1059 (2013). 10.1016/j.cell.2013.07.042

56 Pérez-Verdugo, F., Maniou, E., Galea, G. L. & Banerjee, S. Anisotropic Cell Shape and Motion Coordinate Hindbrain Neuropore Morphogenesis. bioRxiv, 2024.2011.2021.624679 (2025). 10.1101/2024.11.21.624679

57 Mannherz, H. G. et al. Activated cofilin colocalises with Arp2/3 complex in apoptotic blebs during programmed cell death. Eur J Cell Biol 84, 503–515 (2005). 10.1016/j.ejcb.2004.11.008

58 Gagliardi, P. A. et al. MRCKalpha is activated by caspase cleavage to assemble an apical actin ring for epithelial cell extrusion. J Cell Biol 217, 231–249 (2018). 10.1083/jcb.201703044

59 Rasmussen, I. et al. Effects of F/G-actin ratio and actin turn-over rate on NADPH oxidase activity in microglia. BMC Immunol 11, 44 (2010). 10.1186/1471-2172-11-44

60 Niwa, R., Nagata-Ohashi, K., Takeichi, M., Mizuno, K. & Uemura, T. Control of actin reorganization by Slingshot, a family of phosphatases that dephosphorylate ADF/cofilin. Cell 108, 233–246 (2002). 10.1016/s0092-8674(01)00638-9

61 Andrianantoandro, E. & Pollard, T. D. Mechanism of actin filament turnover by severing and nucleation at different concentrations of ADF/cofilin. Mol Cell 24, 13–23 (2006). 10.1016/j.molcel.2006.08.006

62 Breitsprecher, D. et al. Cofilin cooperates with fascin to disassemble filopodial actin filaments. J Cell Sci 124, 3305–3318 (2011). 10.1242/jcs.086934

63 Wang, D. et al. Actin-Depolymerizing Factor and Cofilin-1 Have Unique and Overlapping Functions in Regulating Intestinal Epithelial Junctions and Mucosal Inflammation. Am J Pathol 186, 844–858 (2016). 10.1016/j.ajpath.2015.11.023

64 Sousa-Squiavinato, A. C. M., Rocha, M. R., Barcellos-de-Souza, P., de Souza, W. F. & Morgado-Diaz, J. A. Cofilin-1 signaling mediates epithelial-mesenchymal transition by promoting actin cytoskeleton reorganization and cell-cell adhesion regulation in colorectal cancer cells. Biochim Biophys Acta Mol Cell Res 1866, 418–429 (2019). 10.1016/j.bbamcr.2018.10.003

65 Jodoin, J. N. et al. Stable Force Balance between Epithelial Cells Arises from F-Actin Turnover. Dev Cell 35, 685–697 (2015). 10.1016/j.devcel.2015.11.018

66 Hayakawa, K., Tatsumi, H. & Sokabe, M. Actin filaments function as a tension sensor by tension-dependent binding of cofilin to the filament. J Cell Biol 195, 721–727 (2011). 10.1083/jcb.201102039

67 Humbert, P., Russell, S. & Richardson, H. Dlg, Scribble and Lgl in cell polarity, cell proliferation and cancer. Bioessays 25, 542–553 (2003). 10.1002/bies.10286

68 Jacinto, A. et al. Dynamic actin-based epithelial adhesion and cell matching during Drosophila dorsal closure. Curr Biol 10, 1420–1426 (2000). 10.1016/s0960-9822(00)00796-x

69 Pyrgaki, C., Trainor, P., Hadjantonakis, A. K. & Niswander, L. Dynamic imaging of mammalian neural tube closure. Dev Biol 344, 941–947 (2010). 10.1016/j.ydbio.2010.06.010

70 Abdul-Aziz, N. M., Turmaine, M., Greene, N. D. & Copp, A. J. EphrinA-EphA receptor interactions in mouse spinal neurulation: implications for neural fold fusion. Int J Dev Biol 53, 559–568 (2009). 10.1387/ijdb.082777na

71 Vasioukhin, V., Bauer, C., Yin, M. & Fuchs, E. Directed actin polymerization is the driving force for epithelial cell-cell adhesion. Cell 100, 209–219 (2000). 10.1016/s0092-8674(00)81559-7

72 Hinoi, T. et al. Mouse model of colonic adenoma-carcinoma progression based on somatic Apc inactivation. Cancer Res 67, 9721–9730 (2007). 10.1158/0008-5472.CAN-07-2735

73 Camerer, E. et al. Local protease signaling contributes to neural tube closure in the mouse embryo. Dev Cell 18, 25–38 (2010). 10.1016/j.devcel.2009.11.014

74 Pryor, S. E., Massa, V., Savery, D., Greene, N. D. & Copp, A. J. Convergent extension analysis in mouse whole embryo culture. Methods Mol Biol 839, 133–146 (2012). 10.1007/978-1-61779-510-7_11

75 Marshall, A. R. et al. in *Cell Polarity Signaling*. Methods in Molecular Biology Vol. 2438 (2022).

76 Schindelin, J., et al. Fiji: an open-source platform for biological-image analysis. Nat Methods 9, 676–682 (2012). 10.1038/nmeth.2019

