## Supplementary material for "CFL1-dependency of discrete cell behaviours driving mouse spinal zippering": Figure S

### Supplementary Figures and Legends

### Figure S1: Cell and tissue-level patterns of pCFL1 enrichment. 2

### Figure S2: Surface ectoderm responses to loss of CFL1. 3

### Figure S3: Analysis of lamellipodial and filopodial protrusions. 4

### Figure S4: Pioneer protrusions bridge the embryo midline. 5

Figure S5: Proposed molecular regulation of zippering	6

Supplementary Table 1: Key resource table	7

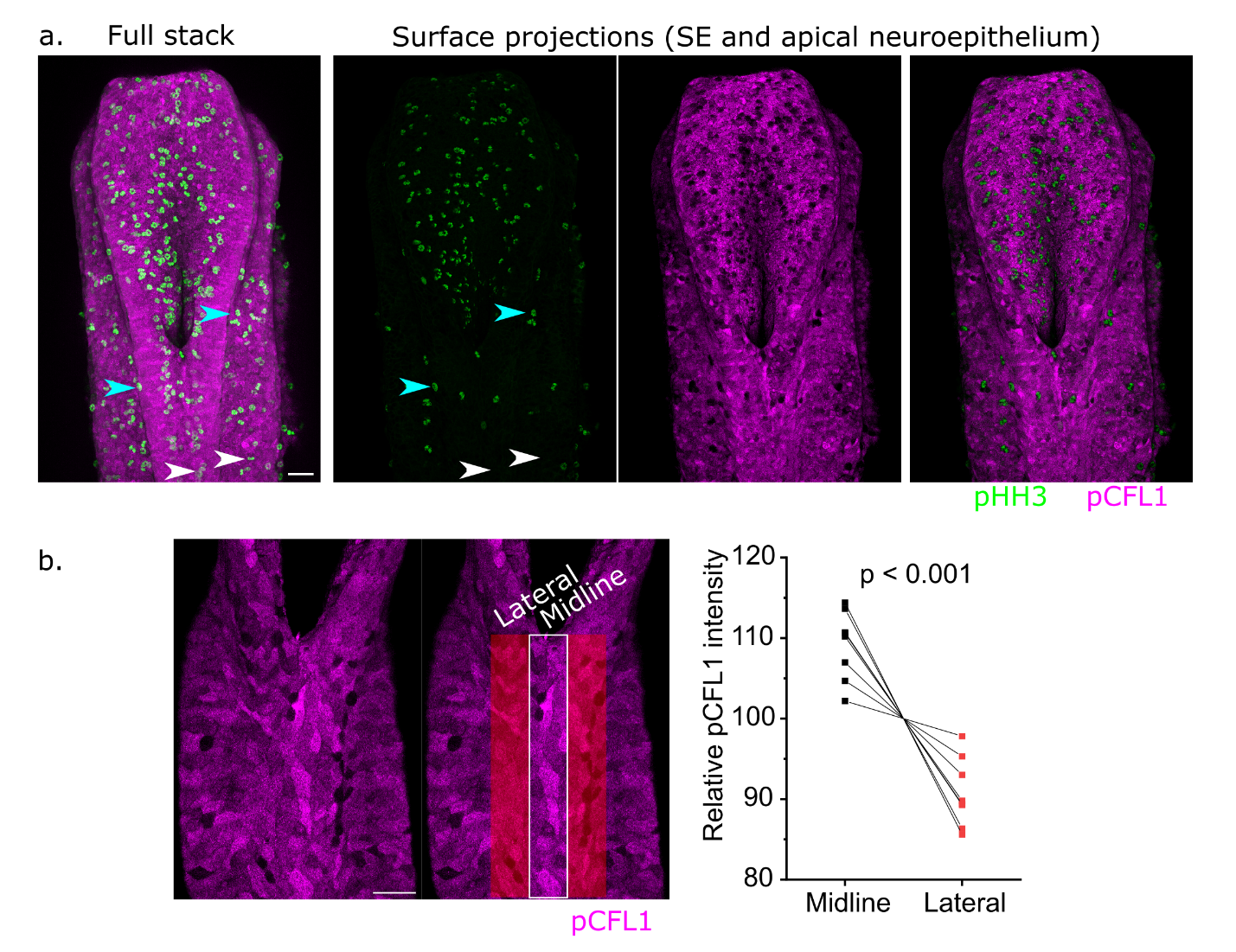

#### Figure S1: Cell and tissue-level patterns of pCFL1 enrichment.

**a.** Illustration of the adaptive surface subtraction method used to isolate dorsal structures including the SE (SE) from a confocal Z-stack of the PNP of an 18 somite stage embryo. The ‘Full stack’ image shows a maximum projection of the Z-stack including both superficial and deep signal, such as from the closed neural tube. The ‘Surface projection’ images only show the top layer of signal across the structure. Note separation of pHH3-labelled mitotic figures in the neuroepithelium of mesoderm (white arrows) from those in the SE (cyan arrows). Also note visualisation of SE pCFL1 independently of the swamping signal from thicker underlying structures.

**b.** Quantification of pCFL1 in a 50 µm band along the embryonic midline versus the lateral regions on either side of the midline. Mean intensity of the two lateral regions was averaged for each embryo and pCFL1 intensity was normalised to 100% across the analysis area of each embryo. P value by paired t-test, points represent individual embryos with their midline and average lateral intensity linked by lines. The illustrative surface subtracted image shows a 16 somite stage embryo.

**a-b.** Scale bars = 50 µm.

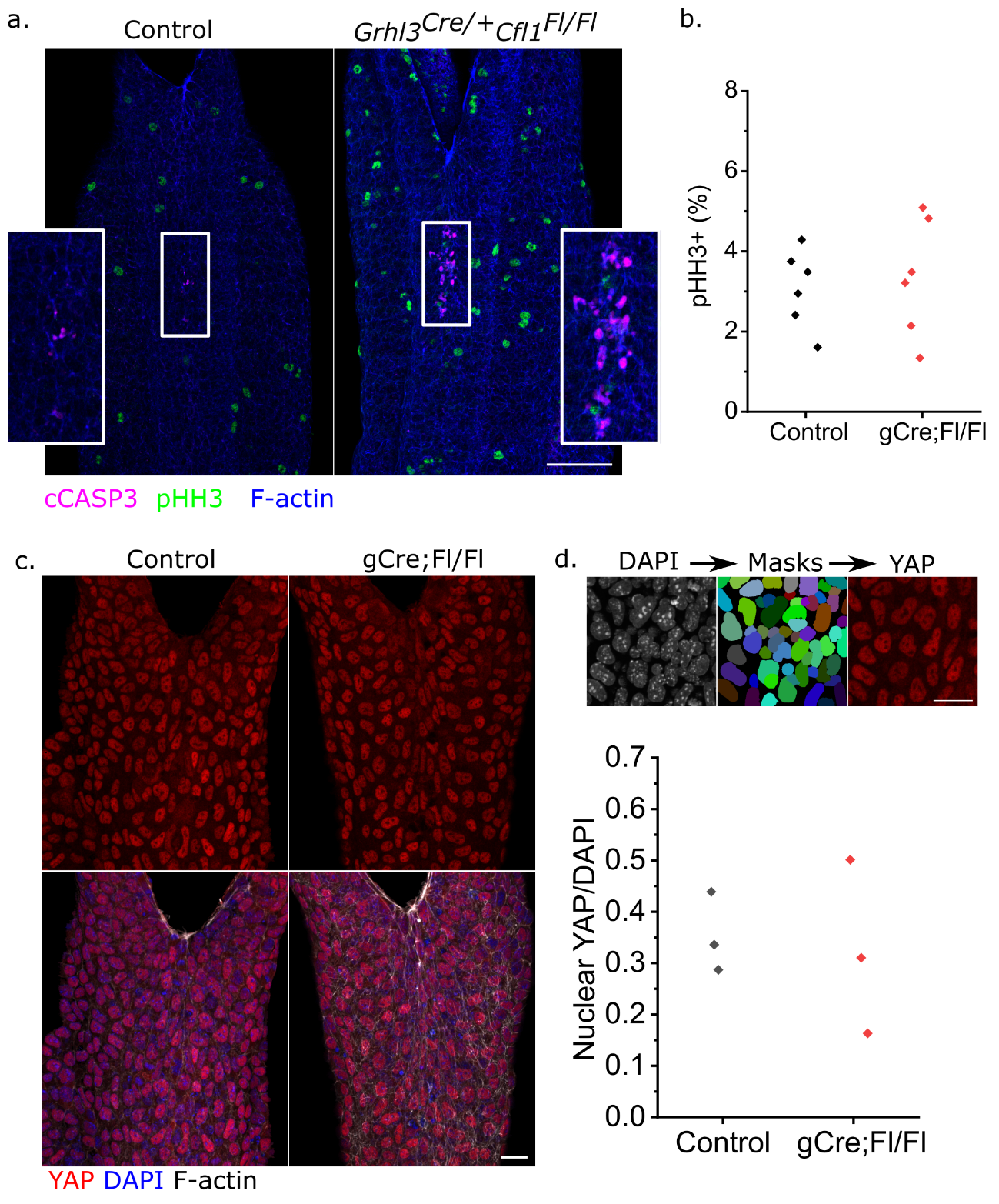

#### Figure S2: Surface ectoderm responses to loss of CFL1.

**a.** Wholemount immunolocalisation of pHH3 and cleaved caspase 3 in the SE of 18 somite control and gCre;Fl/Fl embryos. Inserts show clusters of apoptotic bodies. Scale bar = 100 µm.

**b.** Mitotic index quantification in control and gCre;Fl/Fl embryos. Points represent individual embryos.

**c.** Surface subtracted projection of YAP localisation in the SE of control and gCre;Fl/Fl embryos. Scale bar = 25 µm.

**d.** Illustration of nuclear segmentation used to quantify YAP nuclear intensity, normalised to DAPI intensity within the same nucleus. Points represent individual embryos.

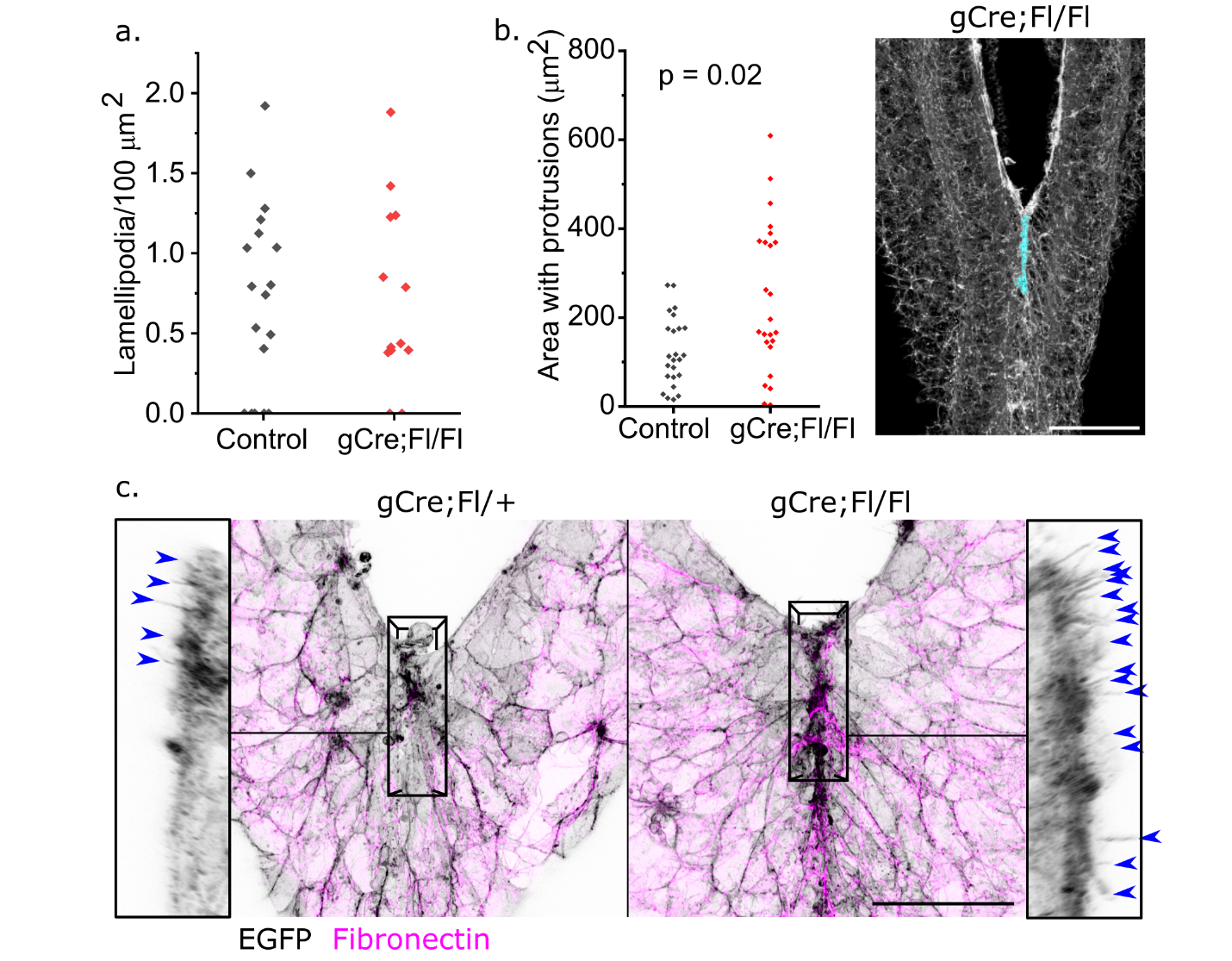

#### Figure S3: Analysis of lamellipodial and filopodial protrusions.

**a.** Quantification of lamellipodial protrusion density around the zippering point in control and gCre;Fl/Fl embryos. Points represent individual embryos.

**b.** Quantification of the area of protrusive activity rostral to the zippering point (schematically illustrated by the cyan colouring in the image of a gCre;Fl/Fl embryo showing F-actin staining, scale bar = 100 µm). Points represent individual embryos.

**c.** Visualisation of fibronectin and surface ectoderm cell membranes lineage-traced with the mTmG reporter in control *Grhl3^Cre/+^Cfl1^Fl/+^Rosa26^mTmG/+^* and littermate *Grhl3^Cre/+^Cfl1^Fl/Fl^Rosa26^mTmG/+^* embryos. The black rectangular outline denotes the region shown in lateral projections adjacent to the main dorsal projection images. Blue arrowheads indicate dorsally-pointing filopodia-like protrusions visualised using the surface ectoderm EGFP lineage trace. Scale bar = 50 µm.

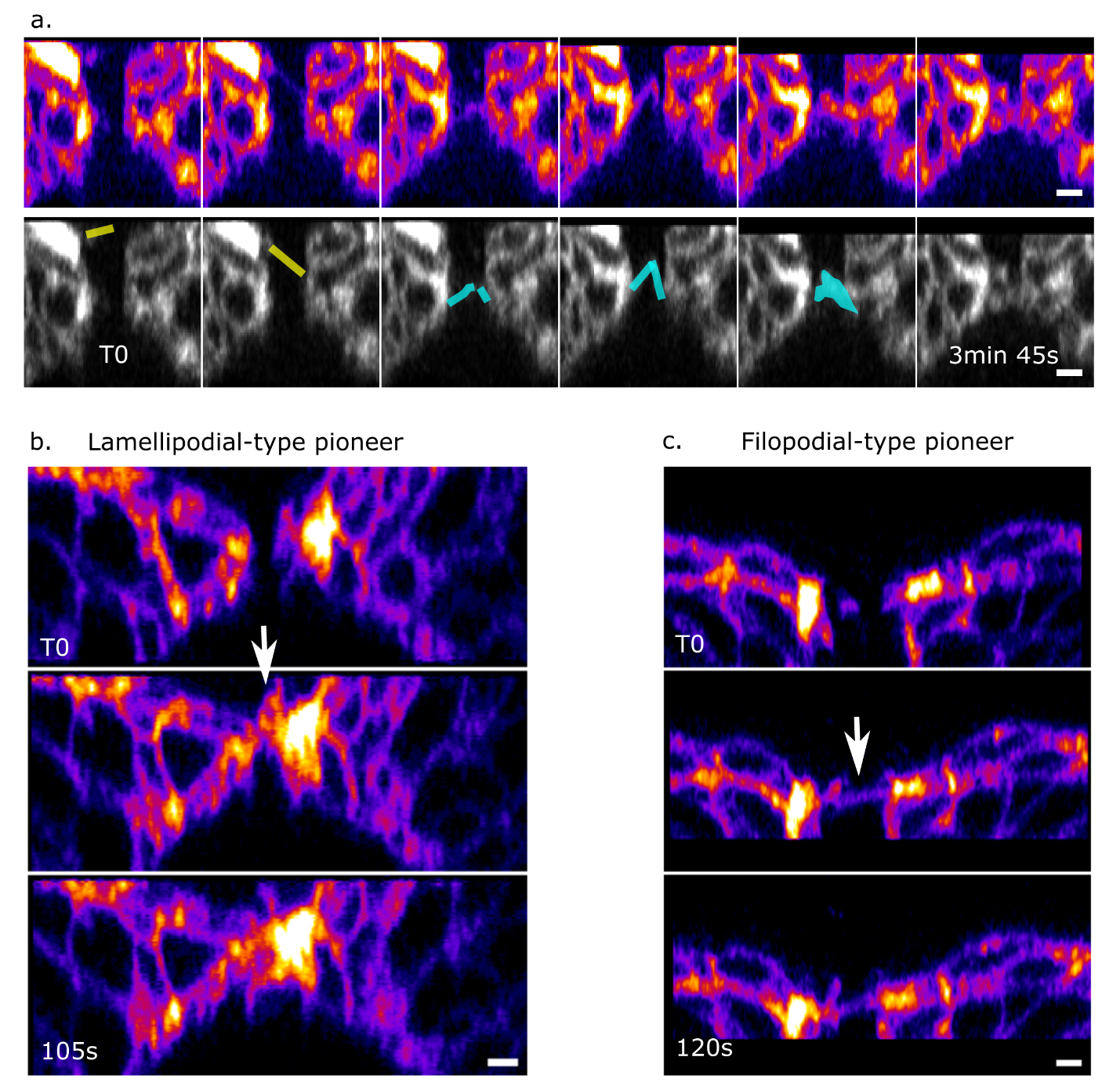

#### Figure S4: Pioneer protrusions bridge the embryo midline.

**a.** Optical cross-section caudal to the zippering point shown in Fire LUT (top) and annotated grey LUT (bottom). Yellow outlines indicate a transient filopodial protrusion which appears to cross the embryonic midline but then retracts. Cyan line indicate pioneer protrusions which meet and are progressively stabilised.

**b-c.** Optical cross-sections showing lamellipodial (**b**) and filopodial (**c**) pioneer protrusions which establish contacts across the embryonic midline and persist.

Scale bars = 5 µm.

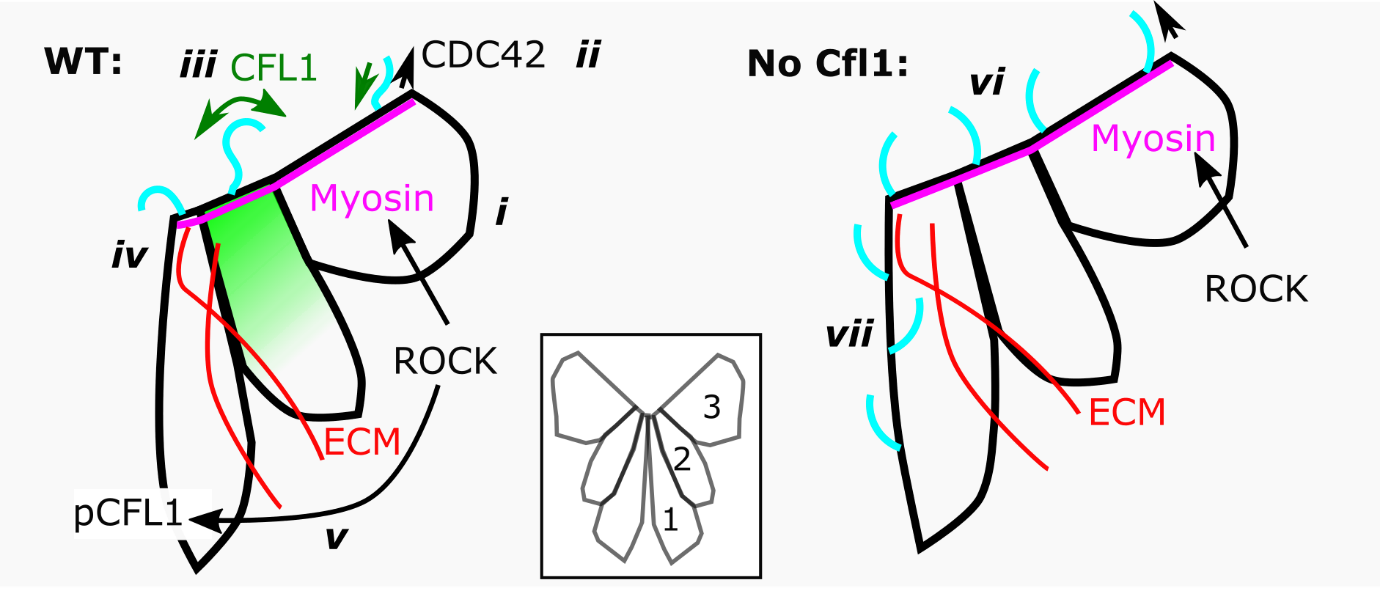

#### Figure S5: Proposed molecular regulation of zippering.

Schematic illustration of the proposed molecular regulation of spinal epithelial fusion by zippering. This schematic combines known molecular mechanisms with findings in the current study on CFL1 action and regulation during mouse spinal zippering. ***i.*** SE cells along the neural fold rim assemble high-tension cables which generate a zippering force through ROCK-dependent myosin contractility^23^. ***ii.*** The same cells simultaneously extend protrusions, including filopodia dependent on CDC42^10^. ***iii.*** CFL1 localises at the leading of the SE cells and in filopodial protrusions, promoting their displacement and timely retraction. ***iv.*** As they approach the zippering point, SE cells undergo CFL1-dependent junctional constriction and form new cell-cell and cell-ECM^19^ contacts. ***v.*** Once no longer connected to the supracellular cables, their ROCK activity promotes phosphorylation of CFL1 instead of cable constriction. ***vi.*** Without CFL1, actomyosin-generated tension is not perturbed and filopodia still extend. ***vii.*** CFL1-depleted filopodia are overly stable, with some persisting rostral to the zippering point.

#### Supplementary Table 1: Key resource table

| **REAGENT or RESOURCE** | **SOURCE** | **IDENTIFIER** |
| --- | --- | --- |
| **Antibodies** | | |
| Rabbit polyclonal anti-CFL1 | Abcam | ab42824 |
| Rabbit monoclonal anti-pCFL1 | Cell Signalling Technology | 3313 |
| Rabbit monoclonal anti-ROCK1 | Abcam | ab45171 |
| Mouse monoclonal anti-pHH3 | Cell Signalling Technology | 9706 |
| Rabbit polyclonal pMLC-II | Cell Signalling Technology | 3671 |
| Mouse monoclonal pMLC-II | Cell Signalling Technology | 3675 |
| Mouse monoclonal E-cadherin | BD Transduction Laboratories | 610181 |
| Rabbit polyclonal anti-cCasp3 | Merck | AB3623 |
| Alexa Fluor 568 Phalloidin | Thermo Fisher Scientific | A12380 |
| Alexa Fluor Plus 405/647 Phalloidin | Thermo Fisher Scientific | A30104 / A30107 |
| **Chemicals, peptides, and recombinant proteins** | | |
| Vybrant^TM^ Multicolor Cell-Labeling Kit | Thermo Fisher Scientific | V22889 |
| Cell Mask Deep Red | Thermo Fisher Scientific | C10046 |
| DirectPCR Lysis Reagent (Mouse Tail) | Viagen Biotech | 102-T |
| Critical commercial assays | | |
| GoTaq G2 Flexi DNA Polymerase kit | Promega | M7806 |
| **Experimental models: Organisms/strains** | | |
| Mouse: wild type C57/BL6 | Bred in-house |  |
| Mouse: Grhl3^tm1(cre)Cgh^ (Grhl3-Cre) |  | MGI: 4430902 |
| Mouse: B6.Cg-Tg(CDX2-cre)101Erf/J (Cdx2-Cre) |  | MGI: 3696953 |
| Mouse: Tg(T-cre/ERT2)1Lwd (T-CreERT2) |  | MGI: 5509185 |
| Mouse: C57BL/6NTac-Tg(ACTB-cre)3Mrt/H |  | MGI: 5316983 |
| Mouse: C57BL/6N-Atm1Brd Cfl1tm1a(KOMP)Mbp/JMmucd | MMRRC | RRID:MMRRC_036512-UCD |
| Mouse: C57BL/6N-Tg(CAG-Flpo)1Afst/Mmucd | MMRRC | RRID:MMRRC_036512-UCD |
| **Oligonucleotides** | | |
| *Cfl1* floxed allele PCR genotyping:  F: GAGATGGCGCAACGCAATTAATG  R: TACTCATAGCACCTCTGAACGTGGC | MMRRC |  |
| *Cfl1* wildtype allele PCR genotyping:  F: ATGACAGGGAGTTGTCCAGCAGG  R: TAGAATGAGTTCCAGGACAGCAGGG | MMRRC |  |
| *Cfl1* null allele PCR genotyping:  F: ATGACAGGGAGTTGTCCAGCAGG  R: TACTCATAGCACCTCTGAACGTGGC | MMRRC |  |
| Cre PCR genotyping:  F: GATGCAACGAGTGATGAGGTTCGC  R: TCCTGGGCAATTTCGGC |  |  |
| **Software and algorithms** | | |
| Origin Lab |  | https://www.originlab.com/ |
| Fiji (ImageJ) |  | https://fiji.sc/ |
| Other | | |
| 11-0 Mersilene microsurgical needles | Ethicon | TG140-6 |
| 10-0 Prolene microsurgical needles | Ethicon | BV75-3 |
